# Cell-wall synthases contribute to bacterial cell-envelope integrity by actively repairing defects

**DOI:** 10.1101/763508

**Authors:** Antoine Vigouroux, Baptiste Cordier, Andrey Aristov, Enno Oldewurtel, Gizem Özbaykal, Thibault Chaze, Mariette Matondo, David Bikard, Sven van Teeffelen

## Abstract

Cell shape and cell-envelope integrity of bacteria are determined by the peptidoglycan cell wall. In rod-shaped *Escherichia coli*, two conserved sets of machinery are essential for cell-wall insertion in the cylindrical part of the cell, the Rod complex and the class-A penicillin-binding proteins (aPBPs). While the Rod complex governs rod-like cell shape, aPBP function is less well understood. aPBPs were previously hypothesized to either work in concert with the Rod complex or to independently repair cell-wall defects. First, we demonstrate through modulation of enzyme levels that class-A PBPs do not contribute to rod-like cell shape but are required for mechanical stability, supporting their independent activity. By combining measurements of cell-wall stiffness, cell-wall insertion, and PBP1b motion at the single-molecule level we then demonstrate that PBP1b, the major class-A PBP, contributes to cell-wall integrity by localizing and inserting peptidoglycan in direct response to local cell-wall defects.

## Introduction

The peptidoglycan cell wall is responsible for both cell shape and mechanical integrity of the bacterial cell envelope (***Typas et al., 2010***; ***Vollmer and Bertsche, 2008***). In Gram-negative bacteria such as *E. coli* the cell wall is a thin two-dimensional polymer that consists of parallel glycan strands oriented circumferentially around the cell axis (***Gan et al., 2008***) and peptide cross-links that connect adjacent glycan strands. To avoid the formation of large pores in the cell wall during growth, cell-wall insertion and cell-wall cleavage must be tightly coordinated (***Vollmer et al., 2008***).

Cell-wall insertion involves two kinds of enzymatic reactions: transglycosylase (TGase) activity to extend the glycan strands, and transpeptidase (TPase) activity to create cross-links between glycan strands. During side-wall elongation these two activities are carried out by two sets of machinery (***Cho et al., 2016***). First, the Rod complex comprises the Penicillin-Binding Protein PBP2, an essential transpeptidase (TPase), and RodA, an essential transglycosylase (TGase) and member of the SEDS (shape, elongation, division and sporulation) family of proteins (***Meeske et al., 2016***; ***Emami et al., 2017***). Together with the MreB cytoskeleton these and other Rod-complex components persistently rotate around the cell (***Lee et al., 2014***; ***Cho et al., 2016***) and are responsible for rod-like cell shape. Second, bi-functional and essential class-A PBPs (aPBP’s) PBP1a and PBP1b carry out both TPase and TGase activities. PBP1a and PBP1b are activated by the outer-membrane lipoprotein cofactors LpoA and LpoB, respectively (***Typas et al., 2010***; ***Paradis-Bleau et al., 2010***; ***Typas et al., 2012***). Mutants in either PBP1a-LpoA or PBP1b-LpoB are viable and don’t show any strong phenotype during regular growth, but mutants in components from both pairs are synthetically lethal (***Yousif et al., 1985***; ***Typas et al., 2010***; ***Paradis-Bleau et al., 2010***). aPBPs also interact with cell-wall cleaving lytic transglycosylases and DD-endopeptidases (***Banzhaf et al., 2019***), consistent with the possibility that they form multi-enzyme complexes responsible for both cell-wall expansion and insertion.

In the past, aPBPs have been suggested to work in close association with the MreB-based Rod complex (***Pazos et al., 2017***), motivated by biochemical interactions between PBP1a and the Rod-complex TPase PBP2 (***Banzhaf et al., 2012***), and by similar interactions between PBP1b and the divisome TPase PBP3 (***Bertsche et al., 2006***). However, each set of enzymes remains active upon inhibition of the respective other one and aPBPs and Rod-complex components show different sub-cellular motion (***Cho et al., 2016***). Furthermore, cells inhibited in PBP1ab activity rapidly lyse (***García del Portillo et al., 1989***; ***Wientjes and Nanninga, 1991***), while cells inhibited in Rod-complex activity become round but don’t lyse (***Lee et al., 2014***).

Since aPBPs and Lpo’s form envelope-spanning complexes (***Egan et al., 2014***; ***Jean et al., 2014***) they have been suggested to work as repair enzymes that activate at site of defects or large pores in the peptidoglycan (***Typas et al., 2012***; ***Cho et al., 2016***). In support of this idea, aPBP activity was increased (***Lai et al., 2017***) upon over-expression of the DD-endopeptidase MepS, which cleaves peptide bonds (***Singh et al., 2012***). Therefore, Rod complex and aPBPs might serve different functions despite catalyzing the same chemical reactions (***Zhao et al., 2017***; ***Pazos et al., 2017***). In agreement with this viewpoint, recent work in the gram-positive *Bacillus subtilis* showed that the two machineries have opposing actions on cell diameter and lead to either circumferentially organized or disordered cell-wall deposition (***Dion et al., 2019***).

Based on the selective interactions between PBP1a-PBP2 and PBP1b-PBP3 (***Banzhaf et al., 2012***; ***Bertsche et al., 2006***), and based on a mild localization of PBP1b at the cell septum (***Bertsche et al., 2006***), PBP1a was suggested to be mostly involved in cell elongation and PBP1b in cell division. However, PBP1b also contributes to cell elongation, where it might have an even more important role than PBP1a under normal growth conditions: PBP1b localizes throughout the cell envelope, with only a mild enrichment at the septum (***Bertsche et al., 2006***; ***Paradis-Bleau et al., 2010***). Furthermore, strains lacking PBP1b have greater mechanical plasticity in the cylindrical part of the cell (***Auer et al., 2016***), their overall rate of peptidoglycan insertion is reduced (***Caparrós et al., 1994***), they are more sensitive to chemicals targeting side-wall elongation, including mecillinam (***García del Portillo and de Pedro, 1991***), A22 (***Nichols et al., 2011***), and D-methionine (***Caparrós et al., 1992***), and they cannot recover from spheroplasts (***Ranjit et al., 2017***).

Here, we study the role of aPBPs for cell shape and cell-wall integrity. First, we measure viability and cell shape during steady-state growth at different protein levels. We found that aPBPs have no role in maintaining cell shape and are therefore not required for proper Rod-complex activity. On the contrary, we confirmed that aPBPs are essential for mechanical cell-wall integrity. Second, we investigate how the major aPBP PBP1b contributes to mechanical integrity: simply through a higher overall rate of peptidoglycan insertion (***Caparrós et al., 1994***), by constitutively stabilizing cell-wall, for example by inserting peptidoglycan in a more spatially homogeneous manner, or through active repair of local cell-wall damage, as previously suggested (***Typas et al., 2012***; ***Lai et al., 2017***; ***Cho et al., 2016***). We first measured mechanical stability and rate of peptidoglycan insertion in cells with aPBP levels reduced three-fold. These cells showed reduced cell-wall stiffness and integrity while maintaining a high rate of peptidoglycan insertion. Therefore, PBP1b apparently strengthens the cell wall independently of changes in insertion rate. Increased integrity could then come about either through constitutive PBP1b activity or through an adaptive repair mechanism (***Typas et al., 2012***). Using a combination of cell-wall perturbations and time-dependent expression of PBP1b, we found that PBP1b facilitates cell survival as quickly as 5 min after protein expression, suggesting that PBP1b senses and repairs cell-wall defects. As a complementary approach, we used single-molecule tracking of a GFP-PBP1b fusion. We found that the bound, non-diffusive fraction of PBP1b molecules decreased with increasing PBP1b or PBP1a levels and increased with LpoB levels, suggesting that PBP1b binds to regions of the cell wall in a need-based manner, which is facilitated through LpoB. Second, we effectively increased the average cell-wall pore size by transiently inhibiting cell-wall insertion during growth. We found that the bound fraction of PBP1b molecules increased shortly after drug treatment and remained high up to 20 min after washout, supporting that PBP1b molecules directly respond to cell-wall damage. Together, our results demonstrate that PBP1b is responsible to maintain the integrity and structural organization of peptidoglycan on a local scale by actively repairing cell-wall defects, while neither of the two aPBPs has a role in cell-shape maintenance.

## Results

### Class-A PBPs are dispensable for cell shape but required for cell-envelope integrity

To investigate the importance of aPBPs for cell shape and cell-wall integrity, we constructed a strain with tunable levels of PBP1a and 1b using partial CRISPR knock-down, which reduces the transcription rate by a fractional amount (***Vigouroux et al., 2018***). To that end, we used the strain LC69 (P_Tet_-dCas9) (***Cui et al., 2018***) and fused PBP1a and PBP1b to RFP (mCherry) and GFP (sfGFP) in their native loci, respectively (strain AV44). We then used combinations of different CRISPR guides targeting GFP and RFP with a variable number of mismatches (***Vigouroux et al., 2018***). To extend the range of possible repression levels, CRISPR guides were expressed in two different forms: i) as a CRISPR RNA (crRNA) co-expressed with the tracrRNA, on the pCRRNAcos vector (***Vigouroux et al., 2018***) or ii) as a single-guide (sgRNA) with fused crRNA and tracrRNA, on the pAV20 vector (***Dion et al., 2019***) (Figure 1A). The CRISPR guides are named according to their complementarity to GFP (G) or RFP (R), Ø designating a control guide. Increasing complementary leads to increased repression (Figure 1 - Supplement 1 and table 8).

To ensure that no truncated or non-fluorescent form of PBP1a or PBP1b was produced, we used bocillin-labeled SDS-page (Figure 1 - Supplement 2A). We quantified PBP1ab protein levels by combining relative mass spectrometry (Data-Independent Acquisition or DIA), absolute mass spectrometry (Parallel Reaction Monitoring or PRM), SDS-page and single-cell fluorescence measurements (see methods and table 1).

**Table 1.** Levels of PBP1ab expressed from different cassettes, and repressed using different sgRNA or crRNA. Ø: Control guides producing no repression. n.d.: not determined. DIA: Data-Independent Acquisition. PRM: Parallel Reaction Monitoring. * Levels relative to LC69 are obtained by multiplying the levels relative to AV44 by the levels obtained by DIA for AV44, with propagated error.

| Relative quantification of PBP1a |  |  |  |  |  |  |
| --- | --- | --- | --- | --- | --- | --- |
| Strain | Promoter | System | Guide | Fluorescence (% of AV44) | Fluorescence (% of LC69*) | DIA (%) |
| LC69 | Wild-type |  |  | n.d. | n.d. | 100 |
| AV44 | Native fusion | sgRNA | R20 | n.d. | n.d. | 20±2 |
| AV44 | Native fusion | crRNA | R20 | 3±4 | 43±56 | n.d. |
| AV44 | Native fusion | crRNA | R18 | 21±4 | 278±139 | n.d. |
| AV44 | Native fusion | crRNA | R11 | 46±6 | 620±298 | n.d. |
| AV44 | Native fusion | crRNA | RØ | 100±7 | 1337±622 | 1337±615 |
| AV63 | P<sub>Bad</sub> | crRNA | R18 | 166±14 | 1549±786 | n.d. |
| AV63 | P<sub>Bad</sub> | crRNA | R11 | 355±22 | 4750±2204 | n.d. |
| AV63 | P<sub>Bad</sub> | crRNA | RØ | 691±50 | 9243±4304 | n.d. |

| Relative and absolute quantification of PBP1b |  |  |  |  |  |  |  |  |
| --- | --- | --- | --- | --- | --- | --- | --- | --- |
| Strain | Promoter | System | Guide | Fluorescence (% of AV44) | Fluorescence (% of LC69*) | DIA (%) | SDS-page (copy/cell) | PRM (copy/cell) |
| LC69 | Wild-type |  |  | n.d. | n.d. | 100 | n.d. | 166±28 |
| AV44 | Native fusion | sgRNA | G20 | 1.0±0.04 | 3.8±0.4 | n.d. | n.d. | n.d. |
| AV44 | Native fusion | sgRNA | G14 | 6.6±0.79 | 24±2.9 | 27±2 | 40±5 | 56±17 |
| AV44 | Native fusion | crRNA | G20 | 4.1±2.0 | 15±7.6 | n.d. | n.d. | n.d. |
| AV44 | Native fusion | crRNA | G14 | 12±3.1 | 44±12 | n.d. | n.d. | n.d. |
| AV44 | Native fusion | crRNA | G10 | 36±2.8 | 131±15 | n.d. | n.d. | n.d. |
| AV44 | Native fusion | crRNA | GØ | 100±5.4 | 367±38 | 367±32 | 688±115 | 547±52 |
| AV58 | P<sub>Bad</sub> | crRNA | GØ | 509±57 | 1870±265 | n.d. | n.d. | n.d. |

**Table 2.** Fit parameters for the mDAP incorporation experiment. The incorporated ^3^H-mDAP per cell is fit with formula 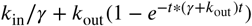, where *k*_in_ is the rate of mDAP incorporation, and *k*_out_ the rate of turn-over, *γ* the growth rate and *t* the time (min). *γ* was measured to be 0.0069 min^-1^, *k*_out_ was fit jointly for all curves and is equal to 0.012 min^-1^. Standard errors for *k*_in_ is also indicated.

| Strain | PBP1a level (%) | PBP1b level (%) | $k_{in}$ (a.u. / min) |
| --- | --- | --- | --- |
| Non-repressed | 1300 | 370 | 786±16 |
| Non-repressed | 1300 | 370 | 943±27 |
| Non-repressed | 1300 | 370 | 886±59 |
| G14-R2 | 20 | 30 | 873±77 |
| G14-R2 | 20 | 30 | 760±67 |
| G14-R2 | 20 | 30 | 863±46 |
| G20-RØ | 1300 | 4 | 678±32 |
| G20-RØ | 1300 | 4 | 706±45 |
| ΔmrcB | 1300 | 0 | 547±51 |
| ΔmrcB | 1300 | 0 | 613±55 |

**Figure 1.**
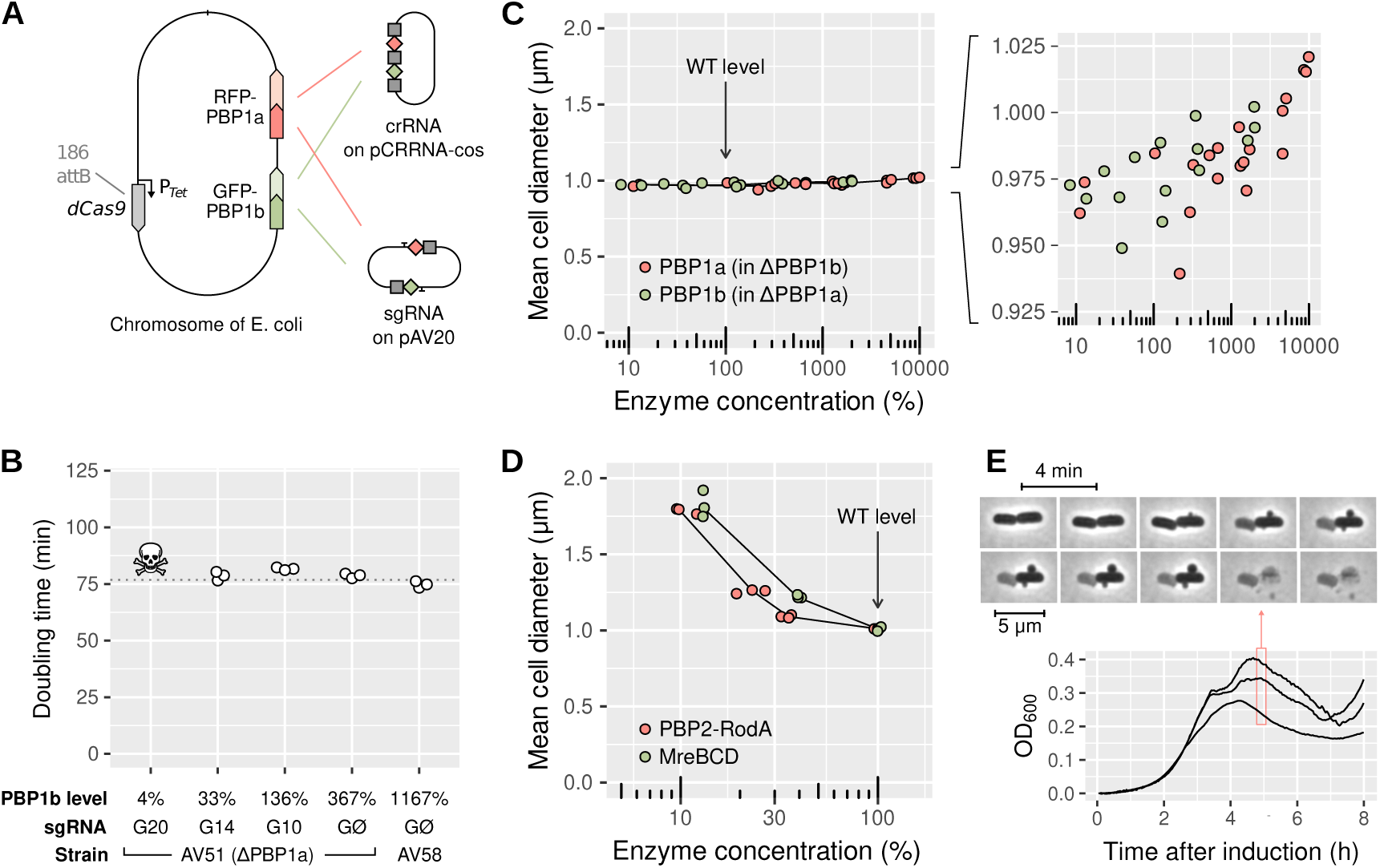
aPBPs have no role in maintaining rod-like cell shape. **A:** Sketch of the strain AV44 (LC69 *mrcB*::*gfp*-*mrcB, mrcA*::*rfp*-*mrcA*) with tunable levels of PBP1a and PBP1b. CRISPR guides are expressed either as crRNA (top) or as sgRNA (bottom), see also Figure 1 - Supplement 1. **B:** Doubling time of AV51 (AV44 ΔPBP1a)/pAV20 as a function of PBP1b level, in minimal medium with glucose and casamino acids at 30°C. sgRNA are expressed from pAV20 as annotated. AV58 is AV51 P_Bad_-GFP-PBP1b for over-expression. Skull logo: not viable. **C:** Effect of aPBP concentration on cell diameter. Points indicate the median diameter within each population. Green: AV51/pCRRNAcos with crRNA G20, G14, G10 and GØ, or AV58 (over-expression). Red: AV50 (AV44 ΔPBP1b)/pCRRNAcos with crRNA R20, R18, R11 and RØ. AV63 is AV50 HK022::P_Bad_-RFP-PBP1a for over-expression. Levels were determined based on fluorescence and normalized with respect to WT according to DIA. **D:** Effect of the concentration of Rod-complex proteins on cell diameter. Green: AV88 (LC69 MreB-msfGFP)/pAV20 with sgRNA G14, G10 or GØ. Red: AV08 (LC69 RFP-PBP2)/pAV20 with crRNA G20, G14, G10 or GØ. **E:** Growth curve of AV44/pAV20 with PBP1ab repressed to lethal level (sgRNA G20-R20), and cell morphology during lysis. Individual points are biological replicates. OD: optical density. WT: wild-type.

The absolute number of PBP1b per cell in the WT is 166±26, in agreement with previous measurements (***Dougherty et al., 1996***). However, levels of non-repressed RFP-PBP1a and GFP-PBP1b were 1300% and 370% higher than their homologs in the wild-type (table 1), reminiscent of previous reports of elevated levels for fluorescent fusions (***Paradis-Bleau et al., 2010***). While not anticipated, this allowed us to explore aPBP levels ranging from strong repression to strong over-expression.

Interestingly, when repressing GFP-PBP1b, the residual expression was higher than the what the same CRISPR guides would produce on constitutive GFP (Figure 1 - Supplement 4, left), suggesting that a form of negative feedback raises PBP1b expression in response to repression. We did not detect such a feedback for RFP-PBP1a (Figure 1 - Supplement 4, right).

As expected from the synthetic lethality of PBP1ab, strains with a strong repression of both PBP1a and PBP1b did not survive. In particular, in strain AV51 (ΔPBP1a), repressing PBP1b to about 4% of WT level using the perfect-match sgRNA G20 (fluorescence microscopy, table 1) leads to cell death. In contrast, AV51 with PBP1b repressed to 30% (DIA, table 1) with sgRNA G14 is still viable. For all the strains which survived repression, the growth rate was unaffected, regardless of aPBPs levels (Figure 1B).

To systematically measure the impact of PBP1ab levels on cell morphology, we varied the level of each PBP between 30 and 1300% (DIA) in strains lacking the respective other PBP. Expression had hardly any effect on cell shape (Figure 1C). From lowest to highest PBP1a or PBP1b levels cell diameter increased by only 75 nm. In contrast, a 10-fold decrease in the level of Rod-complex-related operons PBP2-RodA or MreBCD increased diameter by about 800 nm (Figure 1D), as previously demonstrated (***Vigouroux et al., 2018***). Our observations are also in stark contrast to *B. subtilis*, where a similar change of the level of the major class A PBP PBP1 leads to a 600 nm increase in diameter (***Dion et al., 2019***). As a control, we used an alternative setup based on the inducible P_Bad_ promoter (strains AV100, AV101) and checked the lack of major shape phenotype at low PBP1ab induction (Figure 1 - Supplement 2BC).

We also examined the shape of cells that were depleted for PBP1ab to the point of lysis. To that end we used time-lapse microscopy after induction of our strongest sgRNAs (20 bp of com-plementarity for each target, see table 1). Cells abruptly lysed without changes in cell dimensions compared to the minimum viable expression level (Figures 1E and 1 - Supplement 5). However, we often observed small bulges on the sides of the cells just before lysis. This behavior, previously also observed upon LpoAB depletion (***Typas et al., 2010***), is similar to the effect of beta-lactam antibiotics (***Chung et al., 2009***), suggesting that cells accumulate lethal cell-wall defects in the absence of PBP1ab.

Together, our observations suggest that aPBPs are required for cell-wall integrity at the local scale but dispensable for the maintenance of rod-like cell shape.

### At low levels of aPBPs, cells insert as much peptidoglycan as WT but show reduced mechanical integrity

Next, we aimed to study whether aPBPs maintain cell-wall integrity simply due to an elevated rate of cell-wall insertion or by modulating the cell wall structurally, e.g., through a more homogeneous distribution of peptidoglycan material (***Typas et al., 2012***). It was previously reported that a ΔPBP1b strain inserts about 50% less peptidoglycan, while a ΔPBP1a strain maintains a WT insertion rate (***Caparrós et al., 1994***). We therefore reasoned that the rate of peptidoglycan insertion might not depend on aPBP abundance, as long as a minimum level of PBP1b was present. To study this possibility, we measured the rate of peptidoglycan insertion by recording the incorporation of the radio-labeled cell-wall precursor mDAP (meso-diaminopimelic acid) (***Wientjes et al., 1991***) (Figure 2 - Supplement 1). In AV105 (AV44 ΔPBP1b ΔLysA), peptidoglycan insertion was reduced to about 2/3, even with a high level of PBP1a (Cohen’s d=4.16, p=0.014, t-test, Figure 2A). When repressing PBP1b strongly in AV84 (AV44 ΔLysA) with sgRNA G20, leading to a residual expression of about 4% (table 1), we measured a similar reduction of peptidoglycan insertion as in ΔPBP1b. Intriguingly, when we reduced both PBP1a and PBP1b to about 30% of WT using sgRNA G14-R20 in AV84, the cells inserted

**Figure 2.**
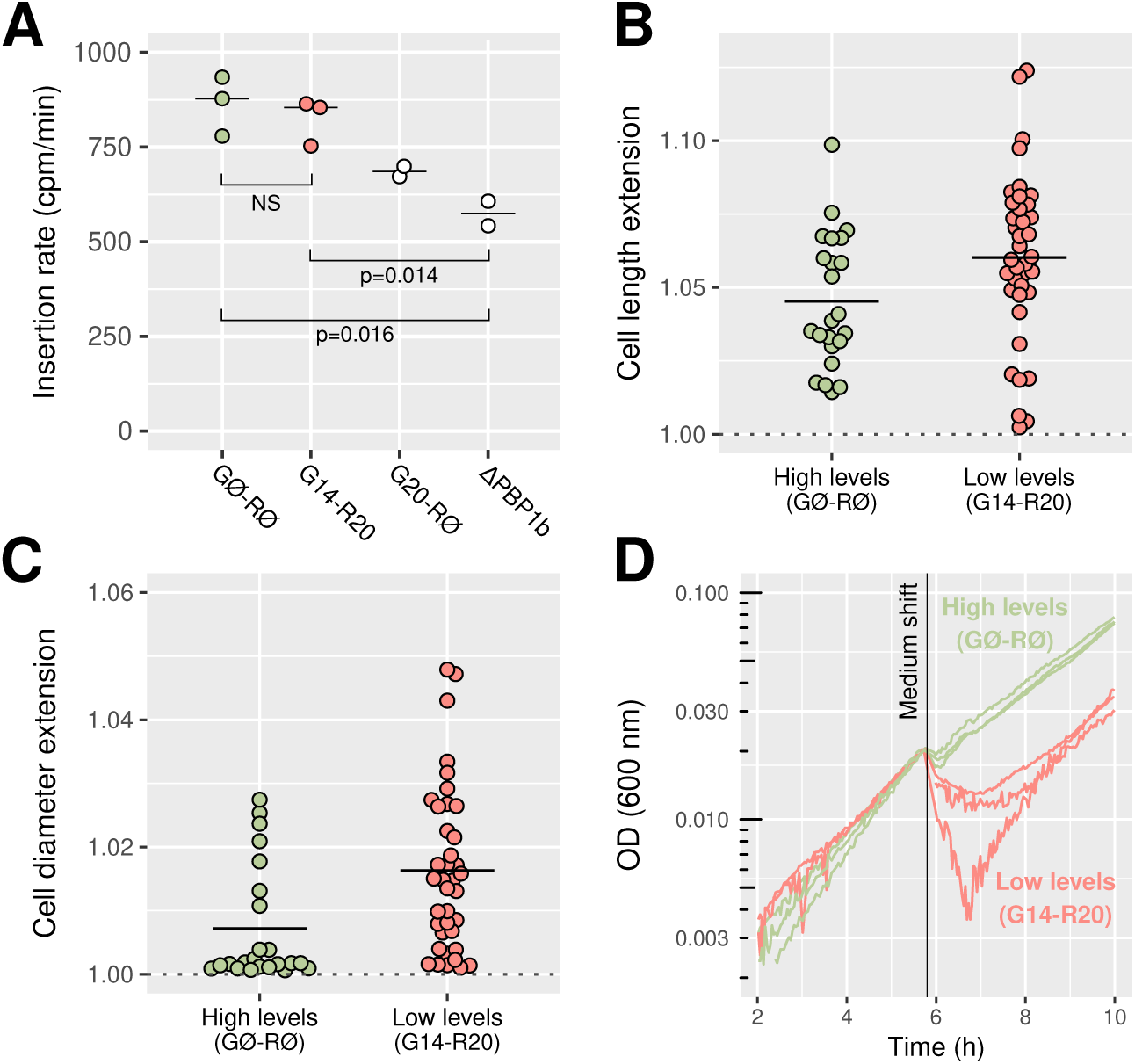
Repression of PBP1ab reduces mechanical stiffness while maintaining a high rate of peptidoglycan insertion. **A:** Rate of insertion of 3H-mDAP into the cell wall measured in AV84 (AV44 Δ*lysA*)/pAV20 and AV105 (AV44 Δ*mrcB* Δ*lysA*) as annotated. NS: not significant. **B-C:** Extension of the cells’ long axis (B) and short axis (C) after a 1 osm/L NaCl downshock, in AV93 (AV44 Δ*mscLS*)/pAV20 with sgRNA GØ-RØ or G14-R20. A value of one corresponds to no extension. **D:** Growth curves before and after a 1 osm/L osmotic downshock, in AV93/pAV20 with sgRNA GØ-RØ or G14-R20. OD: optical density.

PBP1b (Cohen’s d=0.56, p=0.53, t-test, Figure 2A). Therefore, the rate of peptidoglycan insertion is independent of PBP1ab levels as long as PBP1b is expressed at a minimum level between 5-30% of native levels. Furthermore, the decrease of peptidoglycan insertion upon strong PBP1b repression or deletion cannot be compensated by high PBP1a expression, since a strain with 20% of PBP1a and 30% of PBP1b (AV84/pAV20 G14-R20) still inserts more peptidoglycan than a strain with 1300% PBP1a but no PBP1b (AV105/pAV20 GØ-RØ) (Cohen’s d=4.38, p=0.016).

We also measured the chemical composition of the cell wall in these strains through HPLC-UV. No large difference was observed in any of the peaks, meaning that the rate of cross-linking is not affected by the repression (Figure 1 - Supplement 6B), in contrast to what was observed *in vitro* (***Mueller et al., 2019***). Together, these data confirm that peptidoglycan insertion by aPBPs is rigorously buffered against variation in their levels, and buffering holds over a wide range of concentrations.

Next, we wondered whether the reduction of PBPab to low levels might have any effect on the mechanical integrity of the cell wall, even if the rate of peptidoglycan insertion remained high and the composition was unaffected. Previously, it has been reported that cells lacking PBP1b have a more elastic cell wall (***Auer et al., 2016***), a measure of mechanical integrity. To measure potential changes of cell-wall elasticity, we submitted the strain to an osmotic downshock of 1 osm/L of NaCl under the microscope, similarly to (***Buda et al., 2016***) (Fig. 2B and C). To avoid rapid response to osmotic shock, we deleted the mechano-sensitive channels *mscS* and *mscL* from AV44 (strain AV93). The sudden increase of turgor pressure causes an increase of cell dimensions that is inversely related to cell-wall stiffness, in agreement with (***Buda et al., 2016***). We found that repression of PBP1ab to about 30% of WT levels (in AV93/pAV20 G14-R20) leads to a decrease of both axial and circumferential stiffness if compared to the non-repressed strain (Figure 2B and C). Therefore, the reduced number of PBP1ab is likely less capable to protect the cell wall against the accumulation of mechanical defects, despite unperturbed peptidoglycan density and chemical composition.

As a potential consequence of reduced mechanical integrity, we next studied cell survival after osmotic shock in batch culture (Figure 2D and 2 - Supplement 2). In AV93/pAV20, the osmotic shock caused death of a large fraction of cells repressed for PBP1ab (sgRNA G14-R20), while the non-repressed cultures (sgRNA GØ-RØ) were mostly unperturbed.

In summary, we found that cells with reduced levels of PBP1ab showed reduced mechanical stiffness and integrity, which led to an increased rate of cell death upon osmotic downshock, despite unperturbed levels of peptidoglycan density and chemical composition. Therefore, the lack of PBP1ab perturbs cell-wall structure independently of peptidoglycan density.

### PBP1b actively repairs cell-wall damage

PBP1ab could in principle increase mechanical integrity in two different ways: through constitutive peptidoglycan synthesis that compensates the accumulation of mechanical defects but does not respond to the presence of existing defects, or through an active repair mechanism that inserts cell wall in response to damage, as previously proposed (***Typas et al., 2012***). To discriminate these two possibilities, we studied the ability of PBP1b to sustain and recover from transient inhibition of peptidoglycan insertion. Specifically, we blocked peptidoglycan-precursors production by treating cells with the antibiotic D-cycloserine, which inhibits L-alanine to D-alanine conversion and D-alanine-D-alanine ligation (***Lambert and Neuhaus, 1972***), or by starving an auxotrophic mutant strain (*asd-1*) for the essential peptidoglycan component mDAP (***Hatfield et al., 1969***). Different from the above experiments, we expressed PBP1b from a multi-copy plasmid (pBC03) under the control of an inducible P_Bad_ promoter in a ΔPBP1b background for rapid and wide modulation of PBP1b levels. The condition where PBP1b expression is induced will be referred as PBP1b+ and PBP1b-otherwise.

Upon treatment with a high concentration of D-cycloserine (1 mM) under the microscope, cells continued to elongate at a nearly unperturbed rate for about 20-30 minutes before they suddenly lysed (Figure 3A). In batch experiments, WT cells, PBP1b+ cells and PBP1b-cells lysed almost at the same time on average (Figure 3B), demonstrating that the structure of the cell wall prior to drug treatment and the presence of PBP1b during drug treatment have no impact on cell survival.

**Figure 3.**
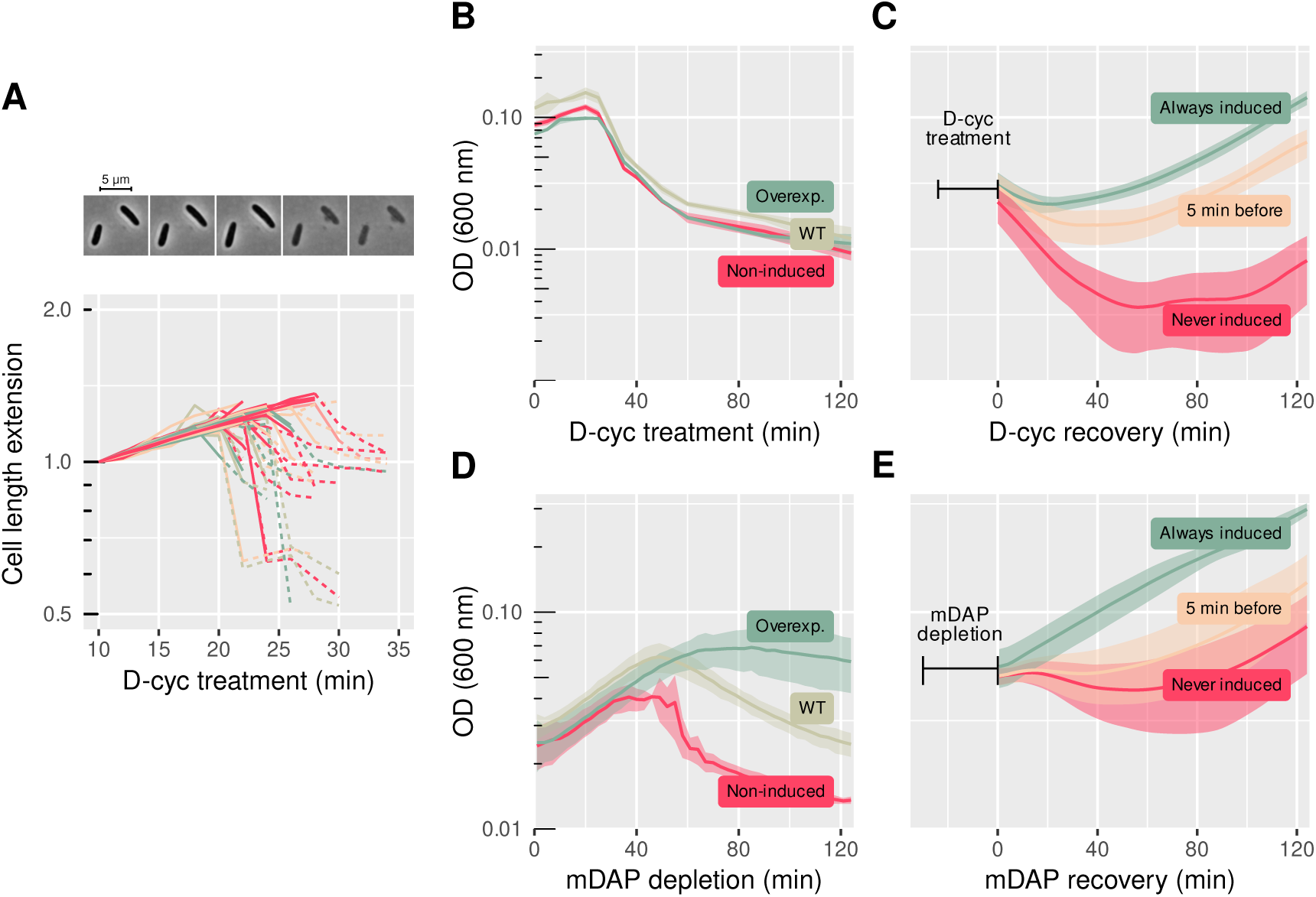
PBP1b facilitates fast recovery from transient inhibition of peptidoglycan synthesis. **A:** Increase of cell length during D-cycloserine treatment (1 mM) under the microscope, with sample snapshots. Strain is AV51 (AV44 ΔPBP1a). Length is normalized by the length at the beginning of the movie. Solid lines describe growing cells, dashed lines correspond to phase-bright, lysing cells. Colors are arbitrary. **B:** Treatment with 1 mM D-cycloserine. Comparison of MG1655 (WT) and B150 (ΔPBP1b)/pBC03 (pBAD33-P_Ara_PBP1b) with arabinose (overexp.) or without arabinose (non-induced). **C:** Recovery of B150/pBC03 after drug washout following 25 min of D-cycloserine treatment (1 mM). PBP1b is either always induced, never induced, or induced 5 min before recovery. **D:** mDAP depletion in the mDAP auxotroph B151 (FB83 *asd-1* (***Teeffelen et al., 2011***)) for WT or B157 (F83 ΔPBP1b *asd-1*)/pBC03 for non-induced/overexp. PBP1b is expressed at different levels like in (C). **E:** Recovery of B157 (F83 ΔPBP1b *asd-1*)/pBC03 after 35 min of mDAP depletion. PBP1b is either always induced, never induced, or induced 5 min before recovery. Shaded areas correspond to mean ± standard deviation of 3 biological replicates. Growth measurements are performed in shaking flasks (B) or microplate reader (C-E). WT: wild-type. OD: optical density.

Notably, cell-wall synthesis was affected well before lysis according to the rotational motion of a fluorescent-protein fusion to MreB (MreB-msfGFP) (***Ouzounov et al., 2016***), which, in turn, requires cell-wall insertion (***Teeffelen et al., 2011***). Within 15 min after drug treatment, processive rotation of MreB-msfGFP stalled (Figure 3 - Supplement 1A), suggesting that cell-wall synthesis was severely reduced at this time. Thus, we reasoned that the density of the cell wall decreased during precursors depletion, as the cells continue to elongate while inserting material at a severely reduced rate.

To study the potential role of PBP1b for cell-wall repair, we washed out D-cycloserine after 25 min of treatment, right before rapid lysis would have started, and monitored growth (Figure 3C). We found that PBP1b-cells showed extensive lysis after drug removal while almost all PBP1b+ cells recovered from damage. To discriminate whether the elevated rate of recovery was due to increased mechanical integrity prior to drug treatment or due to PBP1b activity after washout we compared PBP1b-cells with cells that expressed PBP1b only right before washout. We found that these cells recovered nearly as well as cells expressing PBP1b during the whole experiment. The alleviating effect of PBP1b was almost immediate (<5 min after drug removal). The rapid effect on cell survival suggests that PBP1b actively responds to cell-wall damage and repairs defects.

We observed similar but slightly different behavior upon mDAP depletion and re-addition of mDAP in an mDAP auxotroph (Figure 3D,E). Different from D-cycloserine, mDAP depletion induced lysis only after about 60 min on average in PBP1b+ cells, suggesting that cell-wall synthesis was inhibited later than during D-cycloserine treatment. This is in agreement with previous experiments (***Teeffelen et al., 2011***), where some of us observed a slow reduction of MreB rotation, when cells were grown in minimal medium. Surprisingly, here we observed that MreB-msfGFP rotation was severely reduced within 10-20 min (Figure 3 - Supplement 1C), a similar time as after D-cycloserine treatment. Possibly, a low, non-detected level of cell-wall synthesis was still ongoing.

In contrast to PBP1b+ cells, PBP1b-cells already lysed after about 35 min on average (Figure 3D). We therefore reasoned that PBP1b repaired damage both during and after mDAP depletion.

After re-addition of mDAP following 35 min of mDAP depletion we found that expressing PBP1b right before mDAP repletion had an immediate effect on survival (Figure 3E). However, expressing PBP1b during the whole experiment led to an even faster recovery, presumably because PBP1b helped maintain cell-wall integrity during the 35 min of mDAP depletion.

At a sub-lethal concentrations of D-cycloserine (100 μM), ΔPBP1b cells lysed after about 60 min, while the WT continued to grow, as reported previously (***Nichols et al., 2011***) (Figure 3 - Supplement 2). Similarly to the mDAP depletion experiment, PBP1b+ cells presumably have the capacity to use the reduced pool of peptidoglycan precursors to counter the accumulation of mechanical defects. Together, our findings demonstrate that PBP1b responds to cell-wall damage in an active manner and repairs cell-wall defects.

### PBP1b localizes in response to cell-wall defects

To investigate the active response of PBP1b to cell-wall damage at the molecular level, we studied the movement of individual GFP-PBP1b molecules in the inner membrane. Different from Figure 3, we used minimal medium supplemented with glucose and casamino acids to reduce autofluorescence. Previously, single-molecule tracking of PBP1a in *E. coli* (***Lee et al., 2016***) and PBP1 in *B. subtilis* (***Cho et al., 2016***) revealed that enzymes can be be divided in two populations: a diffusive fraction and a “bound” fraction with near-zero diffusion coefficient. Presumably, only the bound fraction can insert peptidoglycan, while the diffusive fraction is searching for new insertion sites. Notably, bound molecules were detected for a duration of at most a few seconds, which did not allow to identify any persistent motion expected from processive transglycosylation.

To localize individual GFP-PBP1b molecules, we first bleached a large fraction of all molecules in HILO (highly inclined and laminated optical sheet) or epifluorescence mode and then tracked single GFP-PBP1b molecules with an imaging interval of 60 ms in HILO or Total Internal Reflection Fluorescence (TIRF) mode. The fraction of bound molecules was measured by fitting the observed distributions of single-molecule displacements to a two-state model model using the Spot-On tool (***Hansen et al., 2018***) (Figure 4 - Supplement 1). The fit-parameter values for all conditions are indicated in table 3.

**Table 3.** Fit parameters for PBP1b single-molecule tracking. Diffusion constants D_free_ and D_bound_ are in μm^2^/s. The peak localization uncertainty *σ* is in μm. Over: over-expression. WT: approximately wild-type level. Δ: deletion.

| PBP1a | PBP1b | MepS | LpoB | % bound | $D_{\text{free}}$ | $D_{\text{bound}}$ | $\sigma$ | Tracks number |
| --- | --- | --- | --- | --- | --- | --- | --- | --- |
| WT | WT | WT | WT | 18.5 | 0.0681 | 0.007377 | 0.0335 | 8321 |
| WT | WT | WT | WT | 21.1 | 0.0537 | 0.000414 | 0.0376 | 6012 |
| WT | WT | WT | WT | 21.5 | 0.0598 | 0.002609 | 0.0349 | 7156 |

| PBP1a | PBP1b | MepS | LpoB | % bound | $D_{\text{free}}$ | $D_{\text{bound}}$ | $\sigma$ | Tracks number |
| --- | --- | --- | --- | --- | --- | --- | --- | --- |
| 1300% | 30% | WT | WT | 16.88 | 0.0784 | 0 | 0.0437 | 9547 |
| 1300% | 30% | WT | WT | 12.95 | 0.0616 | 0 | 0.0392 | 6986 |
| 1300% | 30% | WT | WT | 18.31 | 0.0751 | 0 | 0.0403 | 7620 |
| $\Delta$ | 370% | WT | WT | 18.03 | 0.0790 | 0 | 0.0547 | 10692 |
| $\Delta$ | 370% | WT | WT | 18.62 | 0.0681 | 0 | 0.0484 | 10806 |
| $\Delta$ | 370% | WT | WT | 20.17 | 0.0768 | 0 | 0.0468 | 9622 |
| $\Delta$ | WT | WT | WT | 20.13 | 0.0864 | 0 | 0.0491 | 13417 |
| $\Delta$ | WT | WT | WT | 23.95 | 0.0721 | 0 | 0.0424 | 7393 |
| $\Delta$ | WT | WT | WT | 25.50 | 0.0752 | 0 | 0.0457 | 10016 |
| $\Delta$ | 30% | WT | WT | 44.57 | 0.0700 | 0 | 0.0430 | 10467 |
| $\Delta$ | 30% | WT | WT | 51.14 | 0.0757 | 0 | 0.0396 | 5371 |
| $\Delta$ | 30% | WT | WT | 41.01 | 0.0722 | 0 | 0.0416 | 10301 |
| WT | WT | WT | $\Delta$ | 7.24 | 0.0579 | 0 | 0.0441 | 9147 |
| WT | WT | WT | $\Delta$ | 10.38 | 0.0717 | 0 | 0.0342 | 7470 |
| WT | WT | WT | $\Delta$ | 16.70 | 0.0622 | 0 | 0.0416 | 10374 |
| WT | WT | WT | Over. | 25.13 | 0.0517 | 0 | 0.0403 | 8280 |
| WT | WT | WT | Over. | 35.96 | 0.0457 | 0 | 0.0336 | 5107 |
| WT | WT | WT | Over. | 34.55 | 0.0450 | 0 | 0.0382 | 7061 |
| WT | WT | WT | WT | 11.00 | 0.0578 | 0 | 0.0373 | 8321 |
| WT | WT | WT | WT | 20.75 | 0.0528 | 0 | 0.0384 | 6012 |
| WT | WT | WT | WT | 19.08 | 0.0543 | 0 | 0.0393 | 7156 |
| 20% | 30% | Over. | WT | 46.94 | 0.0566 | 0 | 0.0380 | 9116 |
| 20% | 30% | Over. | WT | 52.10 | 0.0628 | 0 | 0.0341 | 6137 |
| 20% | 30% | Over. | WT | 40.06 | 0.0701 | 0 | 0.0355 | 8980 |
| 20% | 30% | WT | WT | 34.63 | 0.0623 | 0 | 0.0334 | 6736 |
| 20% | 30% | WT | WT | 41.81 | 0.0706 | 0 | 0.0346 | 7313 |
| 20% | 30% | WT | WT | 38.19 | 0.0626 | 0 | 0.0325 | 7195 |

In order to approximate WT levels in our fluorescently-labeled strain, we used the crRNAs G10 and R18, leading to an expression of 130% for GFP-PBP1b and 280% for RFP-PBP1a (table 1). Using a ΔPBP1a background (AV51) we found about 20% of the PBP1b molecules to be bound around WT levels (crRNA G10), while 80% of the molecules moved diffusively with a diffusion constant of about 0.075 μm^2^/s. Here, bound molecules were found all along the cell axis (and not only at mid-cell) (Figure 4 - Supplement 2).

Qualitatively similar to the observations on the major aPBP PBP1 in *B. subtilis* (***Cho et al., 2016***), we found that the bound fraction of PBP1b decreased with increasing concentration (Figure 4A), suggesting that the activity of individual PBP1b enzymes is reduced upon increasing levels.

**Figure 4.**
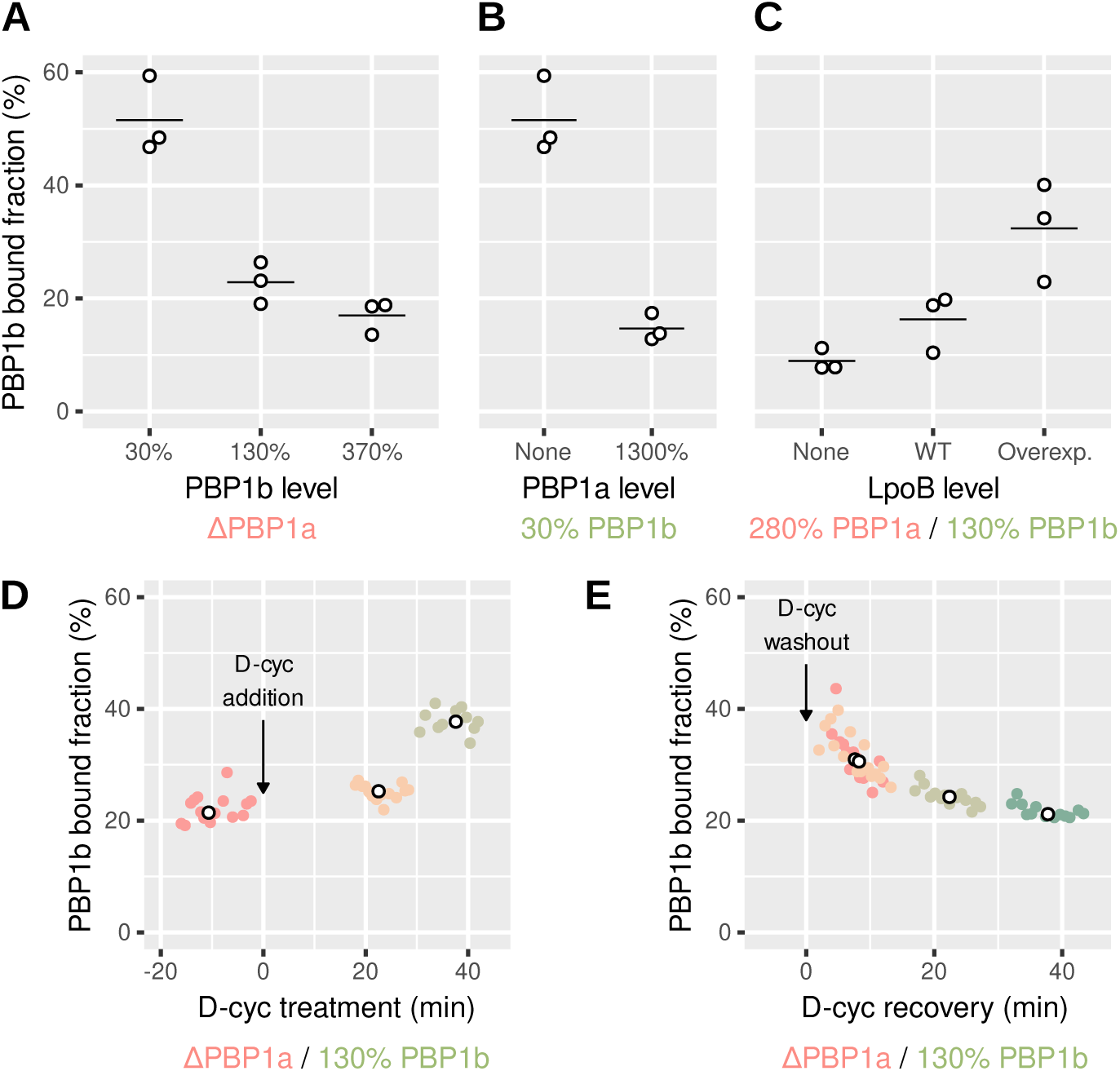
PBP1b localizes depending on the need for peptidoglycan synthesis. **A-D:** Calculated bound fraction of PBP1b at different levels of PBP1b, PBP1a and LpoB, using strains AV44, AV51 (ΔPBP1a) or AV110 (ΔLpoB). For GFP-PBP1b, sgRNA G14 (from pAV20), crRNA G10 (from pCRRNAcos) and crRNA GØ (from pCRRNAcos) are used to reach 30%, 130% and 370% respectively. For RFP-PBP1a, sgRNA R20 (from pAV20), crRNA R18 (from pCRRNAcos) and crRNA RØ (from pCRRNAcos) are used to reach 20%, 278% and 1300% respectively. LpoB is over-expressed using pBC01 (pAM238-P_Lac_-*lpoB*), compared to an empty pAM238 vector, so all cells are grown in the same conditions, in the strain AV44/pCRRNAcos G10-R18. Each point represent a biological replicate comprising at least 5000 tracks. Horizontal lines are means. **D-E:** Bound fraction of PBP1b-sfGFP at different times during D-cycloserine treatment (E) and during recovery from 30 min of D-cycloserine treatment (F) in the strain AV51/pCRRNAcos G10-RØ. Colored points are individual movies and white points are medians for each culture. Corresponding free diffusion coefficients are shown in figure 4- Supplement 4.

The activity of individual PBP1b molecules could be limited by the availability of LpoB molecules, the availability of peptidoglycan precursors, or the abundance of potential sites for cell-wall insertion, henceforth referred to as defects. We therefore aimed to identify the potentially limiting factors by modulating protein levels and precursor availability. First, we modulated LpoB levels. Over-expressing LpoB from plasmid pBC01 (pAM238-P_Lac_-*lpoB*) in AV44/pAV20 G10-R18 indeed increased the bound faction, while deleting LpoB (strain AV110/pAV20 G10-R18) reduced the bound fraction (Figure 4B), indicating that the physical interaction with LpoB aids PBP1b immobilization or stabilizes the bound form.

Next, we investigated how the fraction of bound PBP1b molecules responded to changes of PBP1a abundance. Maintaining PBP1b at about 30% of WT, we found that the bound fraction of PBP1b was reduced about 4-fold upon over-expression of PBP1a (1300% with respect to WT), compared to the ΔPBP1a background (Figure 4C). We reasoned that PBP1a affects PBP1b indirectly through its enzymatic activity, since PBP1a and PBP1b do not share their outer-membrane activators LpoA and LpoB (***Typas et al., 2010***). This could happen through depletion of the common precursor pool or by reducing the number of cell-wall defects detected by PBP1b-LpoB pairs.

To test whether PBP1b immobilization happened immediately after changes of the cell-wall architecture, we transiently inhibited precursor synthesis using D-cycloserine as in figure 3. First, we confirmed that D-cycloserine had the same qualitative effect as in LB (cf. Figure 4 - Supplement 3): Cells lysed after 30-60 min (Figure 4 - Supplement 3A,B), while cell-wall insertion was severely reduced within 10-20 min according to MreB motion (Figure 4 - Supplement 3C,D).

Using single-molecule tracking, we observed a rise of the bound fraction of PBP1b within less than 20 min and a subsequent increase to 37% within 40 min (Figure 4E), while the diffusion constant of diffusive PBP1b molecules was only mildly reduced (Figure 4 - Supplement 4). To make sure that the increase of the bound fraction was due to live, non-lysed cells, we investigated cell shape and also used the nucleic-acid dye propidium iodide, which only penetrates the membranes of dead cells. We then confirmed that visibly dead cells only contributed a small number of tracks to our dataset. Furthermore, while the number of lysed cells visibly rose during the latest set of of movies corresponding to Figure 4 - Supplement 4D (from about 5 to 20% according to visual inspection), we did not observe a concurrent increase of the bound fraction. Our experiments therefore suggest that PBP1b molecules immediately respond to damage by increased binding. Furthermore, the prior arrest of MreB motion suggests that PBP1b binding does not require PBP1b activity or precursor availability.

We then also investigated whether PBP1b showed a higher bound fraction during recovery from D-cyloserine, where the presence of PBP1b greatly increases the chance of cell survival (Figure 3C). In agreement with our expectation, we found that the bound fraction was elevated for about 20 min after a 30 min period of D-cycloserine treatment (Figure 4F).

In summary, our tracking results are compatible with our conclusion above, that PBP1b together with its cognate activator LpoB contributes to cell-wall integrity by localizing and inserting peptidoglycan in response to local cell-wall defects.

## Discussion

In conclusion, we found that different cell-wall-synthesizing machineries have distinct functions in *E. coli*. While the Rod complex is essential for rod shape, the bifunctional aPBPs PBP1ab have hardly any effect on cell shape, up to the point of cell lysis. However, PBP1ab are essential for mechanical cell-wall integrity, and we could demonstrate that PBP1b inserts peptidoglycan in response to local cell-wall defects. Our work therefore contributes to a growing body of evidence suggesting that the local mechanical and structural state of the cell wall provides a major physical cue for peptidoglycan remodeling and insertion.

While cell shape was hardly affected by PBP1ab repression, we found a mild but significant positive correlation between cell diameter PBP1ab levels, consistent with a previous study of a ΔPBP1a mutant (***Banzhaf et al., 2012***). A much stronger correlation between a class-A PBP levels and cell diameter of same sign was recently also observed in *B. subtilis* (***Dion et al., 2019***). In both species, it is conceivable that increased aPBP activity depletes a common pool of lipid II precursors and thus indirectly reduces the capacity of the Rod complex to maintain a narrower cell diameter. What then is responsible for the qualitatively different effect of aPBP levels on cell diameter in *E. coli* and *B. subtilis*? First, our tracking experiments showed that the bound fraction of PBP1b molecules was negatively correlated with PBP1ab expression and positively correlated with LpoB levels. These findings support the model that PBP1b activity is controlled by both the structure of the cell-wall substrate and by the presence of LpoB. In *B. subtilis*, cell-wall synthetic activity of PBP1 might be less regulated, even if PBP1 molecules are less immobile at high expression level (***Cho et al., 2016***). Second, the flux of lipid-II precursors shared by both systems might not be fixed in *E. coli*. Instead, both systems might secure access somewhat independently, as also supported by the overall increase in peptidoglycan synthesis upon MepS over-expression (***Lai et al., 2017***). Finally, LpoB might play an important limiting factor for PBP1b activity, which is absent in *B. subtilis*.

At first sight, our observation of an increasing PBP1b bound fraction with decreasing PBP1a levels seems to be in contradiction to previous measurements of PBP1b diffusion ***Lee et al***. (***2016***). Lee *et al*. reported that deletions of either LpoB or PBP1a hardly affected the average diffusion constant of PBP1b molecules. Similar to our approach, they fused PBP1b to a fluorescent protein (PAmCherry) in the native chromosomal locus. Therefore, it is possible that PBP1b expression was also elevated in their strain, similar to our GFP-PBP1b fusion. We found that at high levels the bound fraction of PBP1b is low, even in the absence of PBP1a (Figure 4A). Expressing PBP1a might then only elicit a small relative change of the PBP1b bound fraction that is hard to detect in the average diffusion constant.

Consistently with average diffusion constants reported in ***Lee et al***. (***2016***) we found that the bound fraction of PBP1b was only mildly reduced upon LpoB deletion, even if both PBP1ab were expressed at native levels. This observation demonstrates that PBP1b does not strictly require LpoB for binding. We reasoned that PBP1b might be able to autonomously detect defects or sites for cell-wall insertion in the absence of LpoB. This hypothesis is consistent with the previous identification of a PBP1b mutant that suppresses the lethality of a ΔPBP1aΔLpoB background (***Markovski et al., 2016***) and with the high residual activity of PBP1b in the absence of LpoB *in vitro* (***Paradis-Bleau et al., 2010***). Alternatively, PBP1b molecules might associate with one or multiple different proteins that immobilizes independently of LpoB. For example, it has been suggested that aPBPs interact with hydrolytic enzymes and with an outer-membrane bound nucleator of cell-wall hydrolases (***Banzhaf et al., 2019***). As another possibility, a fraction of PBP1b molecules might co-localize with the Rod complex or the divisome. We have recently shown that Rod-complex activity remains surprisingly high upon RodA depletion (***Wollrab et al., 2019***) and we reasoned that a different transglycosylase might compensate for the absence of RodA. It will thus be interesting to study the possibility of PBP1b or PBP1a to rescue Rod-complex activity in the absence of RodA in the future.

While repair enzymes are well understood in the context of DNA damage, PBP1b is the first enzyme demonstrated to be involved in the repair of the peptidoglycan cell wall. Yet, the cell wall experiences nearly constant damage due to cell-wall expansion during growth or due to the action of cell-wall antibiotics, making repair all the more important. Recent work by some of us demonstrates that cell-wall cleavage likely happens in regions of elevated mechanical strain and stress (***Wong et al., 2017***). In the absence of repair, increased hydrolytic activity in regions of increased strain would then rapidly lead to more strain and eventually to lysis, as also predicted by computational simulations (***Furchtgott et al., 2011***) and as observed upon depletion of PBP1ab (Figure 1E) or upon treatment with peptidoglycan-synthesis inhibitors (***Yao et al., 2012***). We therefore think that more enzymes might insert peptidoglycan in a manner dependent on the local structure of the cell wall. Consistently, we recently demonstrated that the Rod complex initiates at locations determined by the transpeptidase PBP2, which likely binds to the cell wall directly, in a cell-wall-architecture-dependent manner (***Wollrab et al., 2019***). In the future, the challenge remains to identify the particular local features of the cell wall that attract different cell-wall-modifying enzymes.

## Experimental procedures

### Growth conditions

Cloning and strain preparation were done in Luria-Bertani (LB) medium. Unless mentioned otherwise, every measurement was done in the same M63 with 0.2% glucose, 0.1% casamino acids and 0.5% thiamine. For the experiments of inhibition of peptidoglycan synthesis, cells were grown in LB medium supplemented with arabinose 2 mg/ml if indicated (see below) and mDAP auxotroph strains were grown in LB supplemented with mDAP (50 μg/ml) and L-homoserine (50 μg/ml) from Sigma-Aldrich. For experiments involving the P_bad_ promoter in minimal medium, we used 0.5% lactose instead of glucose as a carbon source. For single-molecule tracking, the concentration of casamino acids used during the preculture and in the agar pad was only 0.01% to minimize background fluorescence.

As needed, media were supplemented with kanamycine (50 μg/ml), carbenicillin (100 μg/ml), chloramphenicol (25 μg/ml) or spectinomycin (50 μg/ml), all from Sigma-Aldrich. CRISPR repression is induced with 100 ng/ml of anhydro-tetracycline (Acros Organics). For over-expression of PBP1a or PBP1b from P_Bad_, 2 mg/ml of arabinose (Sigma-Aldrich) were added to the medium. For over-expression of LpoB or MepS from P_Lac_, we added 1 mM of IPTG. The concentration of propidium iodide used to reveal dead cells was 0.4 μM.

Whenever CRISPR knock-down was employed, dCas9 was induced over night so the repressed gene had time to be diluted to steady-state levels. In the morning, the culture was back-diluted 1/500 and grown for at least 3h to ensure exponential growth before any experiment. Biological replicates result from independent cultures grown from separate colonies.

### Genetic constructions

All strains used in this study derive from the MG1655 and are described in table 4. Plasmids are described in table 5. Gene deletions were carried out starting from the Keio collection (***Baba et al., 2006***). P1 phage lysate was prepared from the Keio deletion strain, then used to infect the recipient strain and the cells were plated on kanamycine to select for transducers. After each phage P1 transduction, as well as all “clonetegrations”, the kanamycine resistance marker was removed with the flippase-expressing pE-FLP (***St-Pierre et al., 2013***). Integration of RFP-PBP1a and GFP-PBP1b in the native locus was done using the allelic exchange procedure described in (***Vigouroux et al., 2018***).

**Table 4.** Strains used in this study.

| Strain | Genotype | Construction |
| --- | --- | --- |
| LC69 | 186::P<sub>Tet75</sub>-*dcas9* | (Cui et al., 2018) |
| AV03 | 186::P<sub>Tet</sub>-*dcas9*, HK022::P<sub>127</sub>, λ::P<sub>127</sub> | Vigouroux et al. (2018) |
| AV04 | 186::P<sub>Tet</sub>-*dcas9*, λ::P<sub>127</sub>-*mcherry* | Vigouroux et al. (2018) |
| AV08 | 186::P<sub>Tet</sub>-*dcas9*, *mrdA*::*mcherry-mrdA* | Vigouroux et al. (2018) |
| AV44 | 186::P<sub>Tet75</sub>-*dcas9*, *mrcB*::*msfgfp-mrcB*, *mrcA*::*mcherry-mrcA* | LC69→pAV42→pAV43 |
| AV47 | 186::P<sub>Tet75</sub>-*dcas9*, HK022::P<sub>127</sub>-*sfgfp*, λ::P<sub>127</sub>-*mcherry* | AV03→P1 (pLC143) |
| AV50 | 186::P<sub>Tet75</sub>-*dcas9*, *mrcA*::*mcherry-mrcA*, Δ*mrcB* | AV44→P1 (Keio Δ*mrcB*) |
| AV51 | 186::P<sub>Tet75</sub>-*dcas9*, *mrcB*::*msfgfp-mrcB*, Δ*mrcA* | AV44→P1 (Keio Δ*mrcA*) |
| AV58 | 186::P<sub>Tet75</sub>-*dcas9*, *mrcB*::*msfgfp-mrcB*, Δ*mrcA*, HK022::P<sub>Bad</sub>-*sfgfp-mrcB* | AV51→pAV71 |
| AV63 | 186::P<sub>Tet75</sub>-*dcas9*, *mrcA*::*mcherry-mrcA*, Δ*mrcB*, HK022::P<sub>Bad</sub>-*mCherry-mrcA* | AV50→pAV77 |
| AV80 | 186::P<sub>Tet75</sub>-*dcas9*, *mrcB*::*msfgfp-mrcB*, *mrcA*::*mcherry-mrcA*, Δ*pbpC*, Δ*mtgA*, Δ*mscS*, Δ*mscL* | AV44→P1 (Keio Δ*mtgA*) |
| AV84 | 186::P<sub>Tet75</sub>-*dcas9*, *mrcB*::*msfgfp-mrcB*, *mrcA*::*mcherry-mrcA*, Δ*pbpC*, Δ*mtgA*, Δ*lysA* | AV80→P1 (Keio Δ*lysA*) |
| AV88 | 186::P<sub>Tet75</sub>-*dcas9*, *mreB*::*mreB-msfGFP* | Dion et al. (2019) |
| AV93 | 186::P<sub>Tet75</sub>-*dcas9*, *mrcB*::*msfgfp-mrcB*, *mrcA*::*mcherry-mrcA*, Δ*pbpC*, Δ*mtgA*, Δ*mscS*, Δ*mscL* | AV80→P1 (Keio Δ*mscS*) |
| AV100 | 186::P<sub>Tet75</sub>-*dcas9*, Δ*mrcA*, Δ*mrcB*, HK022::P<sub>Bad</sub>-*sfgfp-mrcB* | AV58→P1 (Keio Δ*mrcB*) |
| AV101 | 186::P<sub>Tet75</sub>-*dcas9*, Δ*mrcA*, Δ*mrcB*, HK022::P<sub>Bad</sub>-*mcherry-mrcA* | AV63→P1 (Keio Δ*mrcA*) |
| AV105 | 186::P<sub>Tet75</sub>-*dcas9*, Δ*mrcB*, *mrcA*::*mcherry-mrcA*, Δ*pbpC*, Δ*mtgA*, Δ*lysA* | AV93→P1 (Keio Δ*mrcB*) |
| AV109 | 186::P<sub>Tet75</sub>-*dcas9*, *mrcB*::*msfgfp-mrcB*, *mrcA*::*mcherry-mrcA*, Δ*lpoA* | AV44→P1 (Keio Δ*lpoA*) |
| AV110 | 186::P<sub>Tet75</sub>-*dcas9*, *mrcB*::*msfgfp-mrcB*, *mrcA*::*mcherry-mrcA*, Δ*lpoB* | AV44→P1 (Keio Δ*lpoB*) |
| NO34 | *mreB*::*mreB-msfgfp<sub>sw</sub>-kanR* | Ouzounov et al. (2016) |
| B150 | Δ*mrcB* | MG1655→P1 (Keio Δ*mrcB*) |
| B151 | FB83, *asd-1* | Teeffelen et al. (2011) |
| B157 | FB83, *asd-1*, Δ*mrcB* | B151→P1 (Keio Δ*mrcB*) |
| B172 | *mreB*::*mreB-msfgfp<sub>sw</sub>-kanR* | MG1655→P1 (NO34) |
| B174 | Δ*mrcB*, *mreB*::*mreB-msfgfp<sub>sw</sub>-kanR* | B150→P1 (NO34) |
| B176 | FB83, *asd-1*, *mreB*::*mreB-msfgfp<sub>sw</sub>-kanR* | B151→P1 (NO34) |
| B178 | FB83, *asd-1*, Δ*mrcB*, *mreB*::*mreB-msfgfp<sub>sw</sub>-kanR* | B157→P1 (NO34) |

**Table 5.**
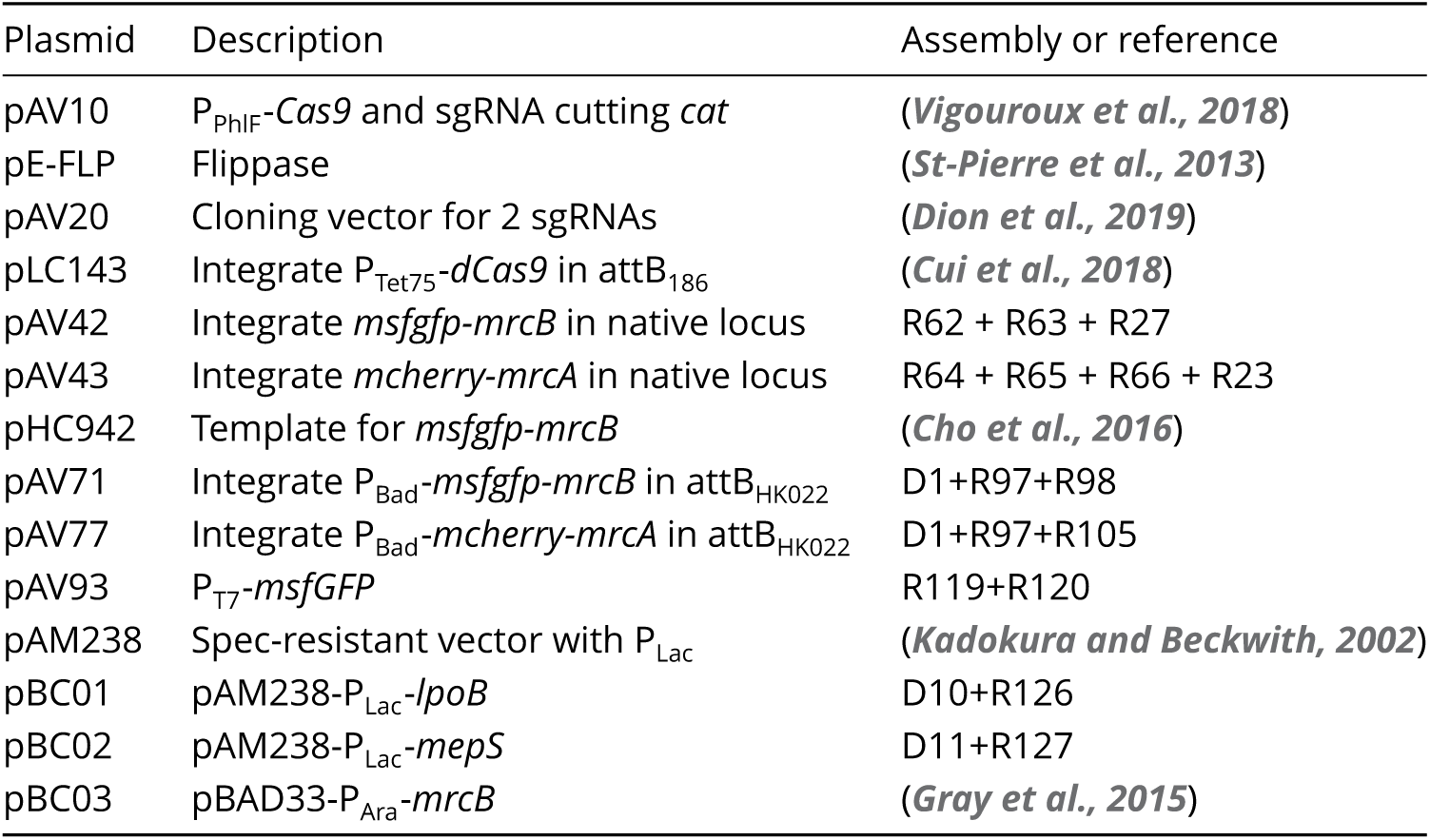
Plasmids used in this study. The fragments “D” are obtained by enzymatic digestions, while the fragments “R” are obtained by PCR amplification. Details about the fragments can be found in table 6. MG1655 is a gift from Didier Mazel.

The plasmids constructed for this study were assembled by Gibson assembly, from the fragments indicated in table 6. Oligonucleotide sequences can be found in table 7.

**Table 6.**
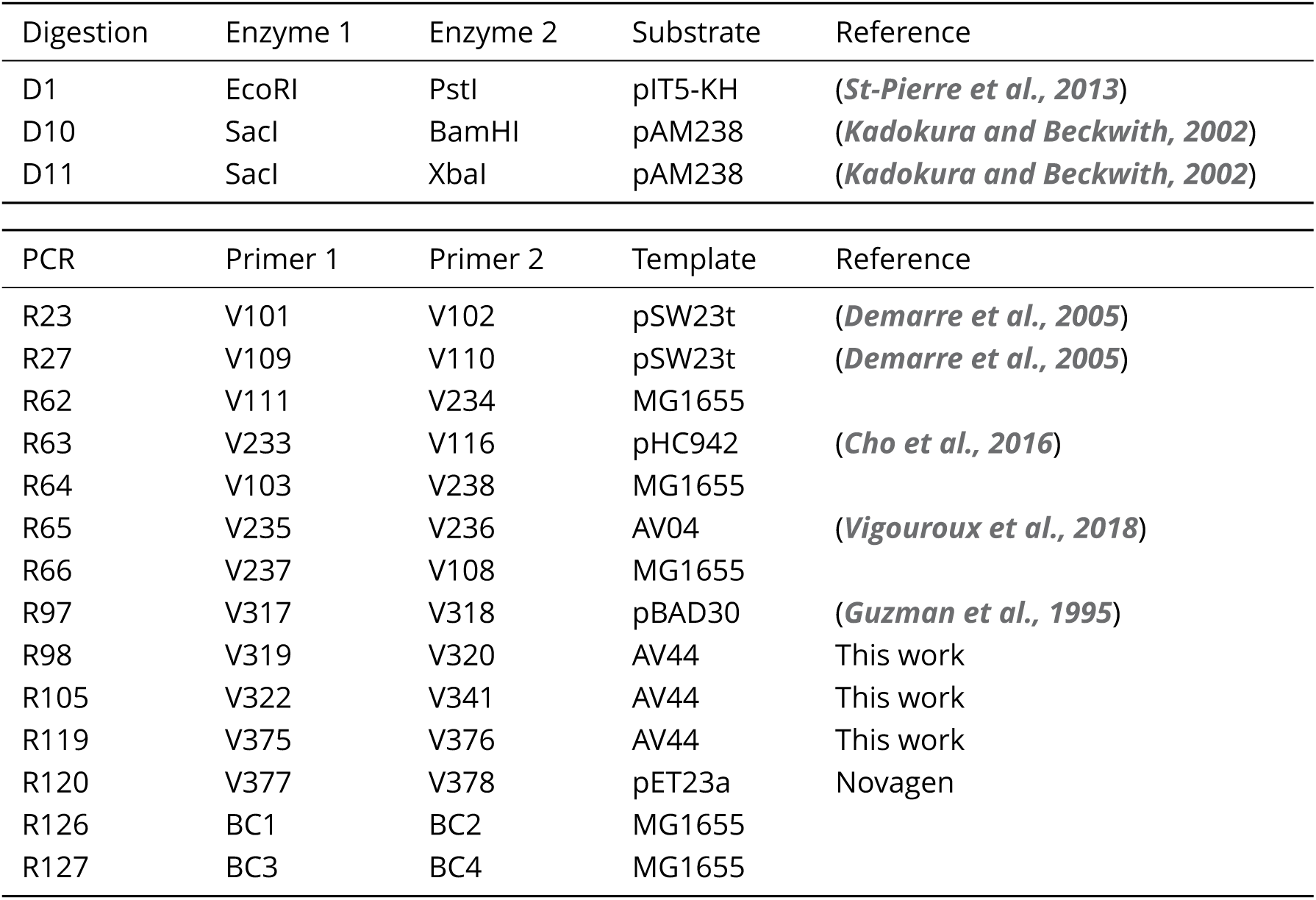
Fragments used to assemble the plasmids.

**Table 7.**
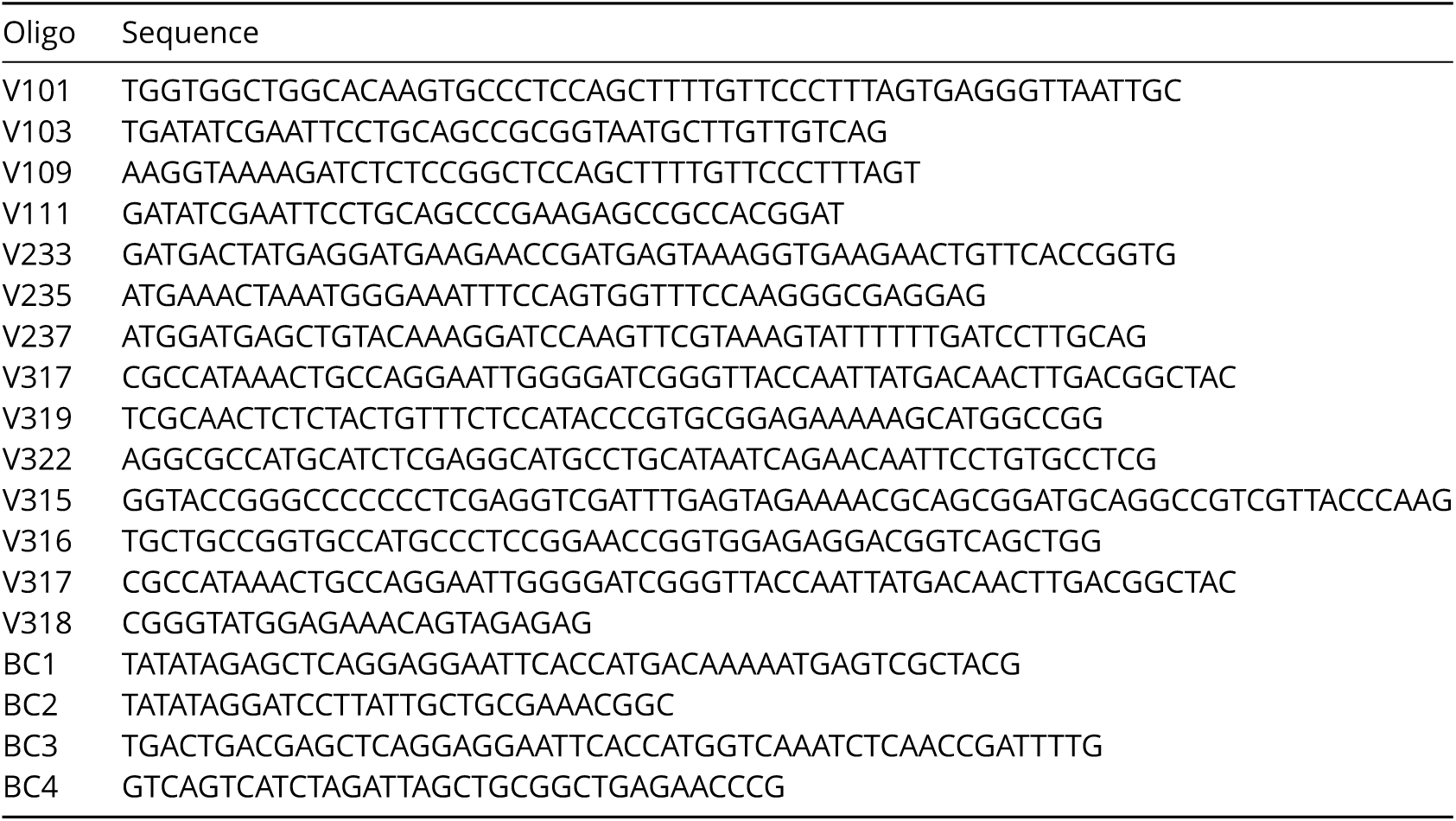
Oligonucleotides used to make the fragments in table 6.

The CRISPR plasmids are either from the pcrRNA collection described in (***Vigouroux et al., 2018***), or were assembled using the pAV20 double-sgRNA vector (***Dion et al., 2019***). In the later case, complementary oligonucleotide pairs (table 8) were phosphorylated with T4 PNK in the presence of T4 ligase buffer (New England Biolabs) and then annealed. A mix containing the pAV20 vector, the two pairs of annealed oligos, the BsaI restriction enzyme (New England Biolabs), T4 ligase (New England Biolabs) and ATP was subjected to thermal cycles for digestion, annealing and ligation. The assembly product was subsequently electroporated in DH5α and the resulting plasmids were sequenced. The “Ø” control guides, producing no repression, still contain the same 5 bp seed sequence as the sfGFP- and RFP-targeting guides. This is to account for potential mild “bad-seed effect” (***Cui et al., 2018***).

**Table 8.**
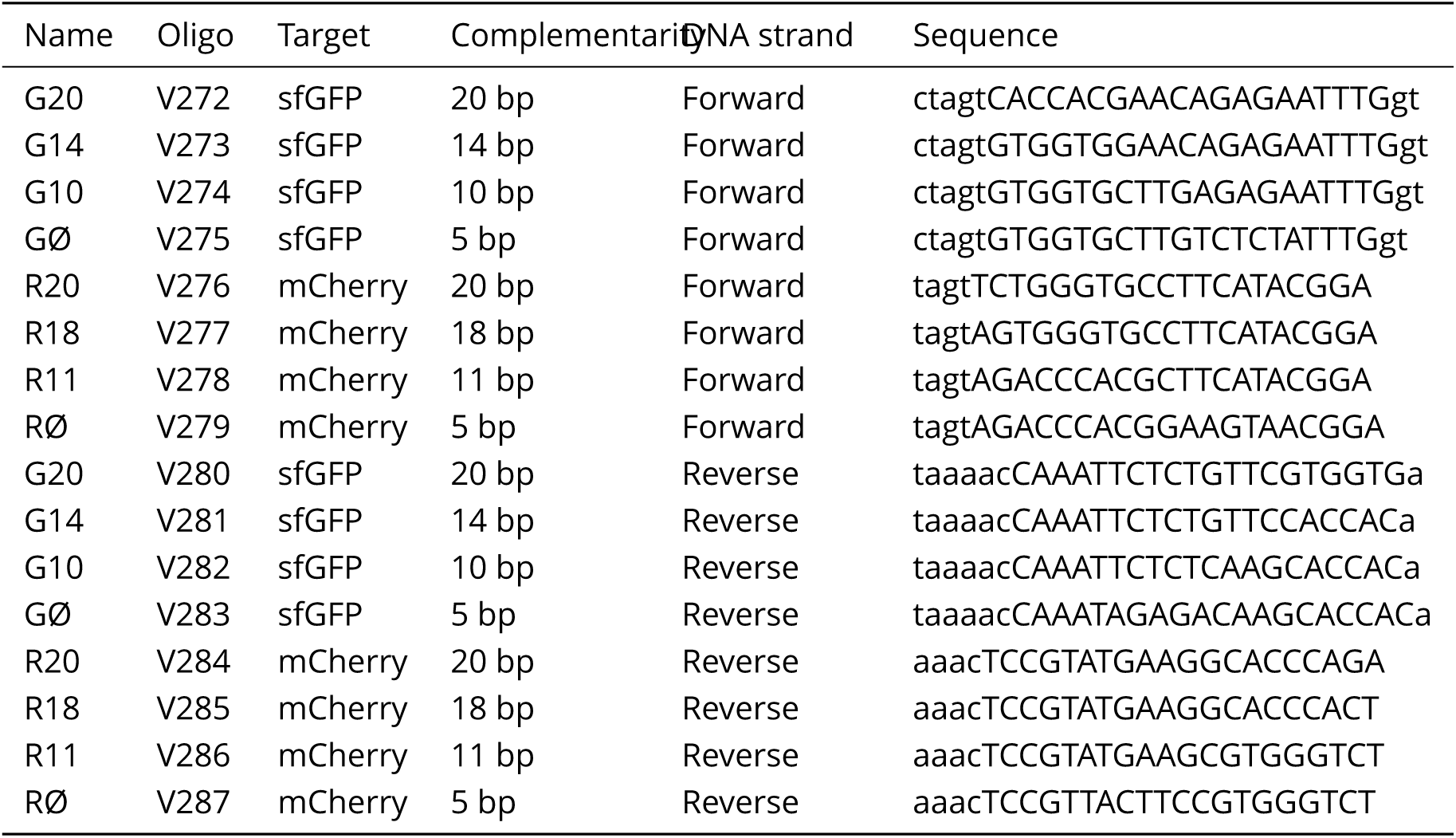
Oligonucleotides inserted in the pAV20 cloning vector to make plasmids expressing two single-guide RNAs, one against GFP (sfGFP) and one against RFP (mCherry). The capital letters indicate the final sequence of the CRISPR guide. After guide insertion, the plasmids are called “pAV20-G*X*-R*Y* “with *X* the complementarity against GFP and *Y* the complementarity against RFP.

### Measurement of optical density and doubling time

Exponential cultures were then transferred to a flat-bottomed 96-microwell plate (Greiner) and optical density at 600 nm was recorded during growth using a microplate reader (Tecan) or, if indicated, using shaking flasks and a spectrophotometer (Eppendorf). To calculate the doubling time, we fit an exponential function to the data points corresponding to the exponential phase. To make sure that the exponential phase was properly isolated, we checked that there was no correlation between consecutive residuals after the fit (Durbin-Watson statistic higher than 1). Optical density at the peak was determined by calculating the first zero of the derivative of OD_600_ after mean-filtering with a bandwidth of 10 min.

### Measurement of cell shape and fluorescence

Cells were grown to steady-state exponential phase (OD_600_ ≈0.1) as detailed in ‘Growth conditions’ and fixed with 4% formaldehyde in phosphate-buffered saline (PBS) for 30 minutes, except for measuring the fluorescence of cells repressed with sgRNA, which were fixed with 1 mg/ml kanamycine in PBS for 30 min. Fixed cells were transferred to agarose pads (1.5% UltraPure Agarose; Invitrogen) containing PBS and imaged using an inverted microscope (TI-E, Nikon Inc.) equipped with a 100× phase-contrast objective (CFI PlanApo LambdaDM100× 1.4NA, Nikon Inc.), a solid-state light source (Spectra X, Lumencor Inc.), a multiband dichroic (69002bs, Chroma Technology Corp.). GFP and RFP fluorescence were measured using excitation filters (560/32 and 485/25 resp.) and emission filters (632/60 and 535/50 resp.). Images were acquired using a sCMOS camera (Orca Flash 4.0, Hamamatsu) with an effective pixel size of 65 nm. The Morphometrics package (***Ursell et al., 2017***) was used to find cell contours from phase-contrast images. Cells that are in proximity from each other were excluded using Morphometrics’ built-in algorithm. In addition, cells were filtered based on their sharpness in phase-contrast (defined as the variance of gradient magnitude). Cell contours were dilated by 1 pixel to capture all the fluorescence of proteins localized to the membrane. For the analysis of fluorescence, we accounted for background intensity, uneven illumination, and cell auto-fluorescence. Intracellular concentration was obtained by integrating the corrected fluorescence intensity inside cell contours, dividing by cell area and subtracting the background value for the image. Total regression was used to find the major axis of the cell. The polar regions were detected by setting a threshold on local contour curvature. Cell width was defined as the average distance between the cell contour and this axis, excluding the poles. Cell length was calculated as the maximal distance between contour points projected on the principal cell axis.

### Quantification of PBP1a and PBP1b

The amount of PBP1a and PBP1b following repression by different CRISPR guides was quantified by several methods.

First, we measured their expression in AV44 pAV20 GØ-RØ (non-repressed), AV44 pAV20 G14-R20 (strong repression) and LC69 (control strain without fusions) using mass spectrometry. We used Data Independent Acquisitions (DIA) (***Bruderer et al., 2017***) for relative quantification of PBP1a and PBP1b. We also used a targeted proteomics approach, Parallel Reaction Monitoring (PRM) (***Bourmaud et al., 2016***; ***Gallien et al., 2012***; ***Peterson et al., 2012***), for absolute quantification of PBP1b. We followed the same protocol previously described (***Wollrab et al., 2019***). Peptides used for absolute quantification of PBP1b were based on the FASTA sequence obtained from UniprotKB database and MS evidence of identification. Peptides sequences are LLEATQYR and TVQGASTLTQQLVK (Aqua UltimateHeavy, Thermo Fisher Scientific).

As a confirmation, we used SDS-page with fluorescence detection to compare AV44 pAV20 GØ-RØ to AV44 pAV20 G14-R20. The detailed procedure is described as supplementary material. Finally, for all the strains whose expression level was not quantified by SDS-page or DIA, we used fluorescence microscopy to mesure relative expression compared to non-repressed AV44, then used the DIA measurement to obtain PBP1ab expression as a percentage of wild-type level. The expression values obtained from the different methods are shown in table 1.

### mDAP incorporation measurement

This experiment was done with the strains AV84 pAV20-GØ-RØ (non-repressed), AV84 pAV20-G14-R20, AV84 pAV20-G20-RØ (ΔPBP1b) and AV105 pAV20-GØ-RØ (20-Ø) (see tables 4 and 8). These strains are lacking *lysA* so radio-labeled mDAP is only used for cell wall synthesis.

Strains were grown to exponential phase and when OD_600_ reached 0.4, ^3^H-labelled mDAP was added for a final activity of 5 μCi/ml. For each time point, 200 μl of culture were transferred to tubes containing 800 μl of boiling 5% SDS. After at least one hour of boiling, the samples were transferred to 0.22 μm GSWP filters. After applying vacuum, the filters were washed twice with 50 ml of hot water. The filters were then moved to 5 ml scintillation vials, treated overnight with 400 μl of 10 mg/ml lysozyme, and dissolved in 5 ml FilterCount cocktail (PerkinElmer) before counting.

The amount of ^3^H-mDAP per cell was calculated by dividing the total counts by the optical density of the culture. The growth rate *γ* was obtained by fitting an exponential function to the OD_600_ values as a function of time. To calculate the incorporation rate *k*_in_ and turn-over rate *k*_out_, we fit the data with formula 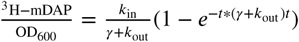 with non-linear least squares optimisation. *k*_out_ is kept constant across all cultures, assuming there are no difference in turn-over rate.

### HPLC content analysis of peptidoglycan

Extraction of peptidoglycan from exponentially growing cells was done according to the protocol described in ***Wheeler et al***. (***2014***). Chromatography of mutanolysin-digested peptidoglycan was performed on a Shimatzu HPLC system with a hypersil Gold eQ 250×4.6 mm column with 3 μm particle size. The mobile phase was a 135 minutes-long gradient from water with 0.05% TFA to 50% acetonitrile with 0.05%TFA. The flow was set to 0.5 ml/min.

### Measurement of cell elasticity and osmotic shock resistance

Osmotic shifts were done by replacing a high-osmolarity medium (M63 with 1/10 volume 5 M NaCl) with a low-osmolarity medium (M63 with 1/10 volume of water) for a shock magnitude of 1 osm. In all cases, cells were grown overnight in high-osmolarity medium then diluted 1/500 and grown at least 3h to reach exponential phase.

To perform osmotic shifts while monitoring cellular dimensions, we constructed a tunnel with two strips of double-sided adhesive tape attached to a glass slide and a cover slip. Polylysine was flushed in the tunnel then washed once with medium. High-osmolarity medium containing a mix of exponentially-growing cells with and without repression of PBP1ab was then flushed in the tunnel. The slide was incubated 15 min for cells to settle. Fresh medium was flushed again to remove unattached cells. Then we took images of GFP and RFP fluorescence, to quantify the total amount of GFP-PBP1b and RFP-PBP1a in each cell. This allowed to distinguish non-repressed cells from repressed ones without ambiguity. We finally recorded phase-contrast images while the low-osmolarity medium was flushed into the tunnel. Cells were tracked using a simple nearest-neighbor algorithm, discarding the cells that went out of focus.

For osmotic shock resistance, cells were prepared in a similar manner, then growth in high-osmolarity was monitored for 6 h in a plate reader. The plate was then centrifuged 2 min at 2000 x *g*, medium was discarded and low-osmolarity medium was added instead. Growth was then monitored again for 4 h.

### Measurement of mreB-msfGFP motion

Fluorescence images were generated on an inverted epi-fluorescence microscope (Nikon Ti-E) equipped with a 100x phase contrast objective (CFI PlanApo LambdaDM100X 1.4NA, Nikon), a solid-state light source (Spectra X, Lumencor Inc., Beaverton, OR), a multiband dichroic (69002bs, Chroma Technology Corp., Bellows Falls, VT), and excitation (485/25) and emission (535/50) filters. Images were acquired using a sCMOS camera (Orca Flash 4.0, Hamamatsu, Japan) with an effective pixel size of 65 nm. For measurements of cell boundaries, we focused on cells based on the phase-contrast signal. To track MreB-msfGFP spots moving at the bottom of the cell, we moved the focal plane 250 nm below the central plane of cells. Images were taken every 2 s for a duration of 120 s.

Images were analyzed using a custom Matlab code as described previously described (***Wollrab et al., 2019***). Briefly, images were first filtered in both space and time using a three-dimensional Savitzky-Golay filter with a filter size of 3 pixels in xy-directions time 3 points along the temporal dimension. Images were subsequently de-noised once more using a 2D-Gauss filter (*σ* = 0.5 pixels). Images were subsequently rescaled by a factor of 5 using spline interpolation to achieve sub-pixel resolution. MreB spots were detected as local maxima inside the cell boundary obtained by segmentation using the Morphometrics package (***Ursell et al., 2017***). MreB spots with intensity higher than the cell background were considered for tracking. The local maxima were connected to construct raw trajectories based on their distance at consecutive time points (***Teeffelen et al., 2011***) with a maximal displacement during subsequent time frames of 3 pixels. After generating the tracks, we applied a Gauss filter in time (*σ* = 1.5 time steps) in order to decrease spatial noise. Tracks which have more than 7 localizations were considered for velocity distributions. Velocity is calculated from single displacement vectors of the smoothened trajectories. Flux is then calculated by summing over all end-to-end distances of smoothened tracks that are longer than 200 nm, normalized by total duration of the movie (2 minutes for all movies) and total surface area of all cells.

### Single-particle tracking of PBP1b

Single particle tracking of sfGFP-PBP1b was performed in either of two custom-designed fluorescence microscopes, equipped with a custom-built temperature controlled chamber at 29°C or a stage-top incubation chamber (Okolab). Prior to imaging cells were transferred to a pre-heated 1% agarose pad (Invitrogen) and covered with a pre-cleaned cover slip. Cover slips were cleaned by bath sonication in a 1M KOH solution for 1h at 40C. Both microscopes were equipped with a 100x TIRF objective (Apo TIRF, 100x, NA 1.49, Nikon), three laser lines: 405 nm (Obis, Coherent), 488 nm (Sapphire, Coherent), 561 nm (Sapphire, Coherent), a dichroic beamsplitter (Di03-R488/561-t3-25×36, Semrock) and a laser-line filter (NF561-18, Thorlabs). Shuttering of the 488 nm laser was controlled with an acousto-optic tunable filter (AA Optoelectronics) or with shutters (Uniblitz, LS3 and TS6B, Vincent Associates). Images were acquired with an EMCCD camera (iXon Ultra, Andor). All components were controlled and synchronized using MicroManager (***Edelstein et al., 2010***). Images were acquired with exposure time and intervals of 60 ms for a duration of 20 s to 1 min. To distinguish single molecules, this requires a photobleaching phase prior to image acquisition. To that end, the sample was exposed to 488 nm laser in epifluorescence or HILO (highly inclined and laminated optical sheet) mode. Since both modalities resulted in the same fractions of bound molecules, the bleaching or illumination modality did apparently not bias towards either state of molecules. After photobleaching, we either switched to TIR mode or remained in HILO mode for image acquisition. Bleaching time is adjusted according to the level of PBP1b and illumination intensity and was about 2 or 12 s for HILO and epi illumination, respectively. A longer bleaching time of 10 or 25 s was required for GFP-PBP1b overexpression (Figure 4A, AV51 without repression).

To determine the bound fraction of GFP-PBP1b, we first segmented images using the brightfield channel and standard image processing functions. PBP1b spots in fluorescence images were identified using the ThunderStorm plug-in for ImageJ (***Ovesný et al., 2014***) with wavelet filtering. The peak detection threshold was equal to the standard deviation of the first wavelet levels of input image (Wave.F1). Sub-pixel resolution was achieved by finding the center of a two-dimensional Gaussian fitted to the intensity profile of each spot. Spots in subsequent frames were then connected using the nearest-neighbor algorithm from TrackPy with a maximum step length of 500 nm (***Allan et al., 2016***). To limit tracking mistakes, we discarded the frames where the peak density was too high by only taking the last 30,000 peaks of each movie into account. The displacements were fit using a two-state diffusion model from the SpotOn software package (***Hansen et al., 2018***), allowing to recover the percentage of bound molecules, the peak localization precision and the free molecules’ diffusion constant. In the reference strain (strain AV44 with near-WT levels of PBP1a and PBP1b) the diffusion constant of the “bound” molecules was left as a free parameter and found to be compatible with immobilization of these molecules (*D*_bound_ *<* 0.001 μm^2^/s, table 3, top). For the rest of the analysis we fixed *D*_bound_ =0 (table 3, bottom).

### Bocillin-labeling of the PBPs

The bocillin-binding assay used to check the absence of non-fluorescent PBP1ab is similar to what is used in (***Cho et al., 2016***; ***Kocaoglu et al., 2012***). We prepared exponentially-growing cells at OD_600_≈0.4. We washed 1.8 ml of each culture in PBS, resuspended them in 200 μl PBS and kept cultures on ice. We disrupted cells by sonication (FB120, Fisher Scientific) and centrifuged them for 15 min at 4°C (21,000 g). We subsequently resuspended the pellet corresponding to the membrane fraction in 50 μl PBS containing 15 μM fluorescently labelled Bocillin-FL (Invitrogen). Membranes were incubated at 37°C for 30 min and washed once in 1 ml PBS. We centrifuged the membranes for 15 min (21,000 g) and resuspended them in 50 μl PBS to remove unbound Bocillin-FL. We measured the protein concentration of each sample with a colorimetric assay based on the Bradford method (Bio-Rad) and loaded equal amounts of protein mixed with 4X Laemmli buffer onto a 10% polyacrylamide gel (Miniprotean TGX, Bio-rad). We visualized the labelled proteins with a Typhoon 9000 FLA imager (GE Healthcare) with excitation at 488 nm and emission at 530 nm.

## Acknowledgements

We thank T. Bernhardt for providing the pHC942 plasmid and W. Vollmer for providing the pBAD33-pbp1b plasmid. We thank Richard Wheeler for assistance with peptidoglycan extraction and chromatography analysis and to Eva Wollrab for help in the single-molecule tracking. This work was supported by the European Research Council (ERC) under the Europe Union’s Horizon 2020 research and innovation program [Grant Agreement No. (679980)], the French Government’s Investissement d’Avenir program Laboratoire d’Excellence “Integrative Biology of Emerging Infectious Diseases” (ANR-10-LABX-62-IBEID), the Mairie de Paris “Emergence(s)” program, and the Volkswagen Foundation.

---

**Figure 1 - Supplement 1.**
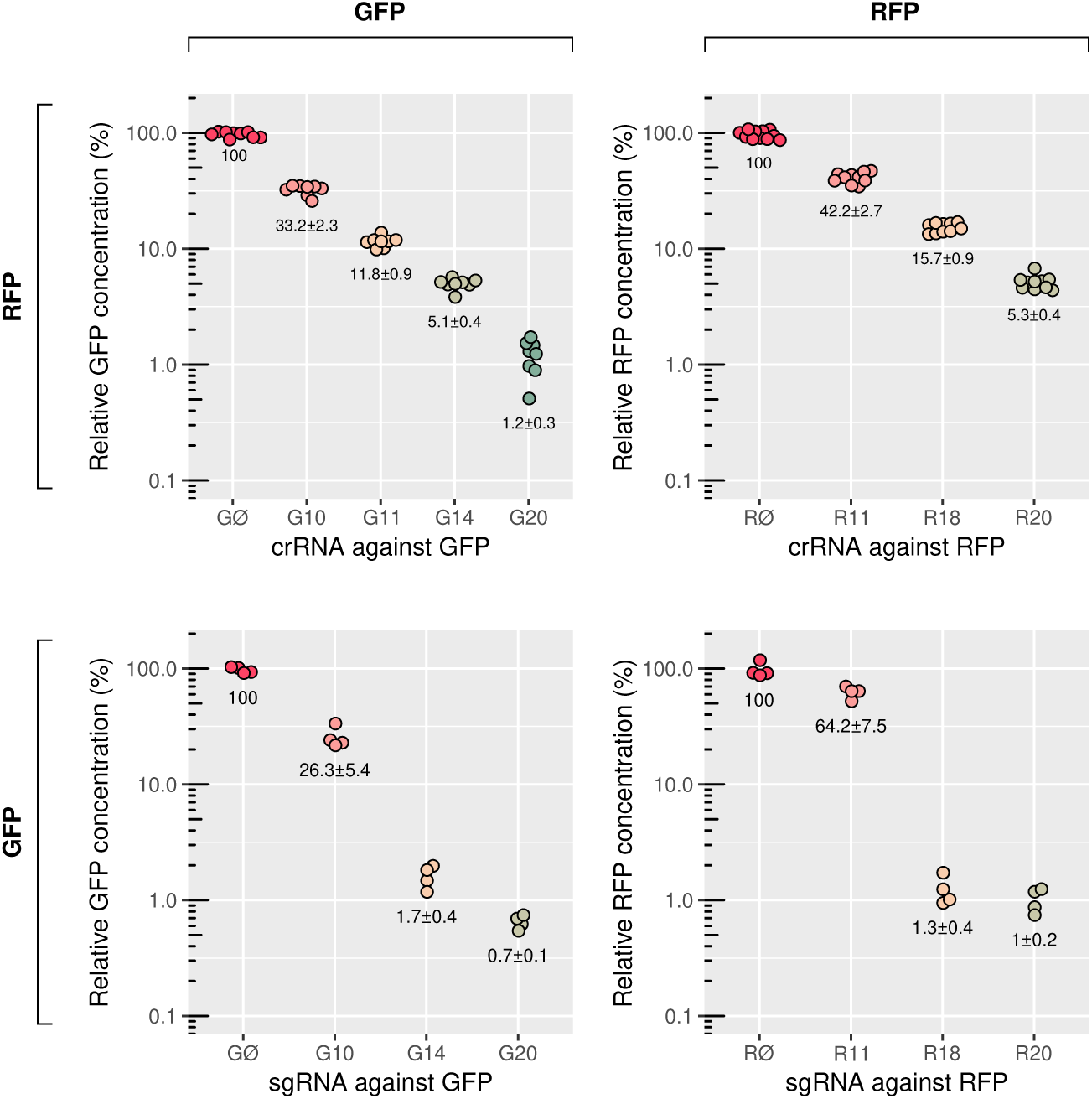
Passage probability of the different CRISPR guides used in this study. This is measured by fluorescence microscopy in strain AV47 (LC69 HK022::P_127_-*sfgfp*, λ::P_127_-*mcherry*)/pCRRNAcos for crRNA or pAV20 for sgRNA. When used to repress the RFP-PBP1a and GFP-PBP1b fusions in AV44, the repression level may be different because of genetic feedback. **A:** Repression of GFP and RFP by crRNAs expressed from the pcrRNAcos vector. **B:** Repression of GFP and RFP by sgRNAs expressed from the pAV20 vector. See table 8 for the sequences of the guides. Relative fluorescence is expressed as a percentage of AV47/pAV20 GØ-RØ, i.e. without repression.

**Figure 1 - Supplement 2.**
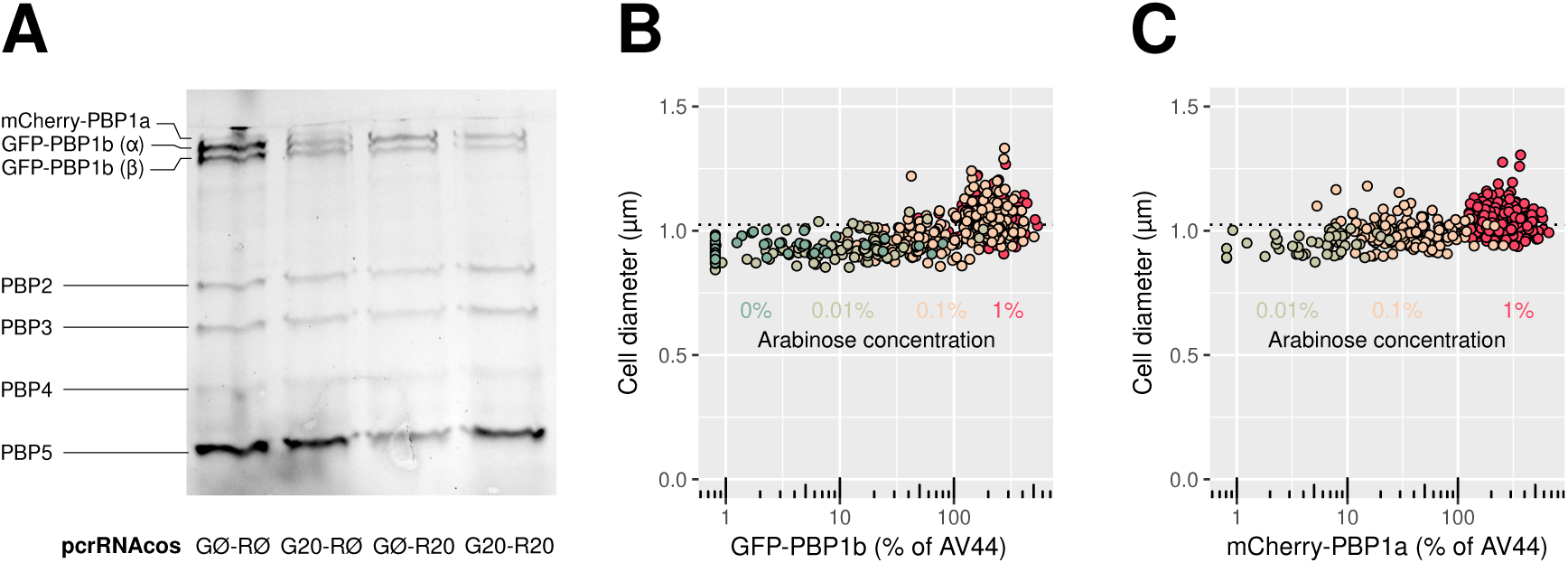
**A:** The RFP-PBP1a and GFP-PBP1b fusions are the only forms of aPBPs present in AV44. Fluorescent bocillin binds specifically to Penicillin Binding Proteins (PBP). The change in band intensity after repression by CRISPR does not reflect the change in fluorescence measured by microscopy for the same conditions, presumably because bocillin only labels potentially active molecules. All experiments are done in AV44/pcrRNAcos with crRNA as annotated. **B and C:** Diameter of single cells at different levels of GFP-PBP1b (B) or RFP-PBP1a fusion (C), in strains AV100 (AV51 HK022::P_Bad_-GFP-PBP1b) and AV101 (AV50 HK022::P_Bad_-RFP-PBP1a) respectively. Different colors indicate different concentrations of arabinose, from 0% to 1%.

**Figure 1 - Supplement 3.**
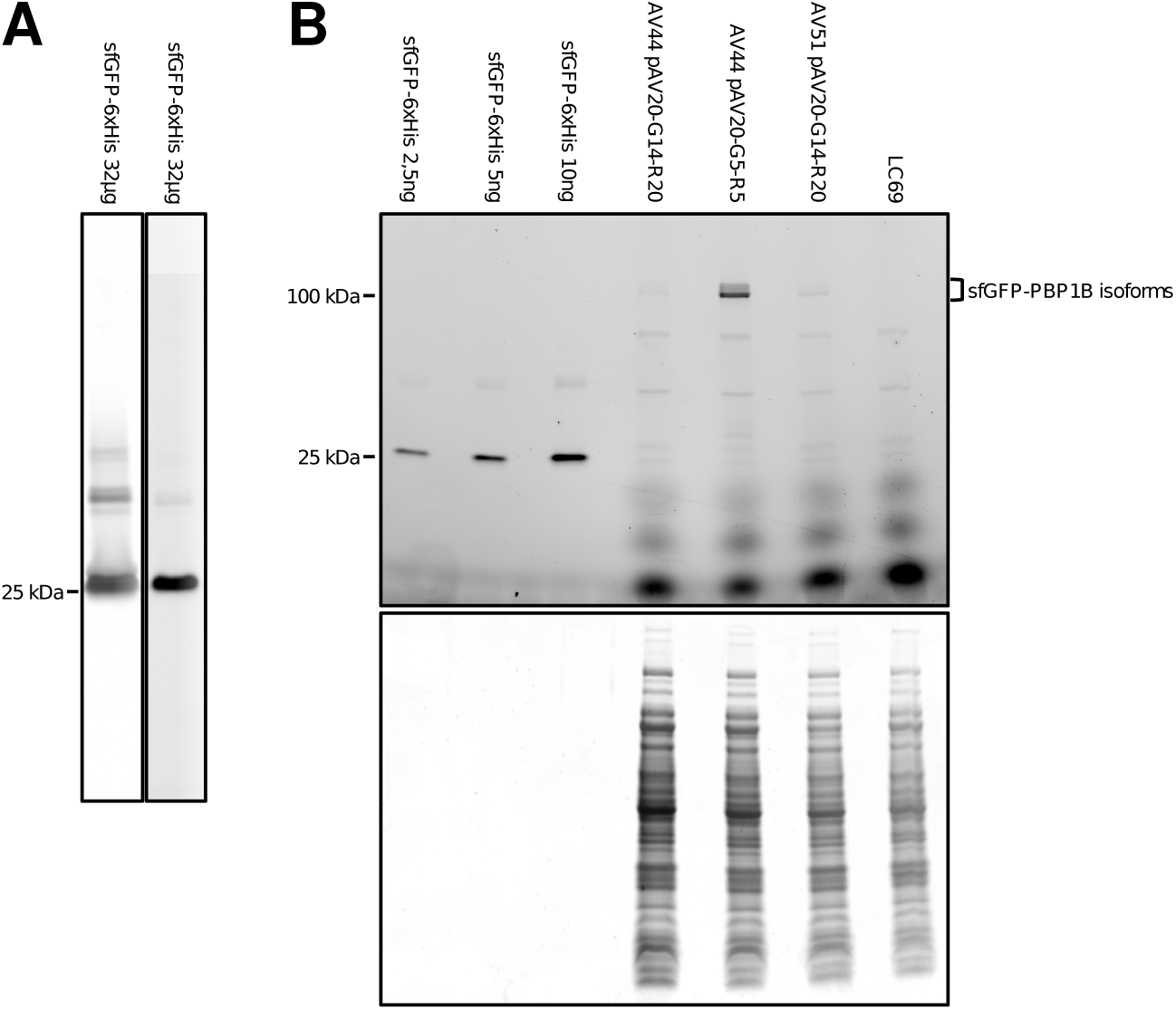
Quantification of sfGFP-PBP1b by semi-quantitative SDS-page. **A:** Purified sfGFP-6xHis. Left: Elution fraction loaded in a 4-20% acrylamide gel stained with Coomassie blue. The predicted sfGFP-6xHis molecular weight is ≈28,42 kDa. Right: visualization of the in-gel fluorescent signal. The higher molecular weight bands probably correspond to sfGFP oligomers as they are also detected on purified sfGFP. **B:** 4-20% acrylamide gel with decreasing amounts of purified sfGFP-6xHis (first 3 lanes), followed with whole cell extracts of LC69, AV44 or AV51 with pAV20 as annotated. Approximately 30 μg of proteins were loaded. sfGFP-PBP1b isoforms: isoform α (predicted molecular weight 94,32 kDa, 121,1 kDa tagged with sfGFP), isoform μ (predicted molecular weight 88,91 kDa, 115,69 kDa tagged with sfGFP). Top: GFP fluorescence measurement. Bottom: Coomassie blue staining. There is no signal of the purified sfGFP-6xHis fusion protein because the amount loaded is probably below visualization limit with the Coomassie blue method.

**Figure 1 - Supplement 4.**
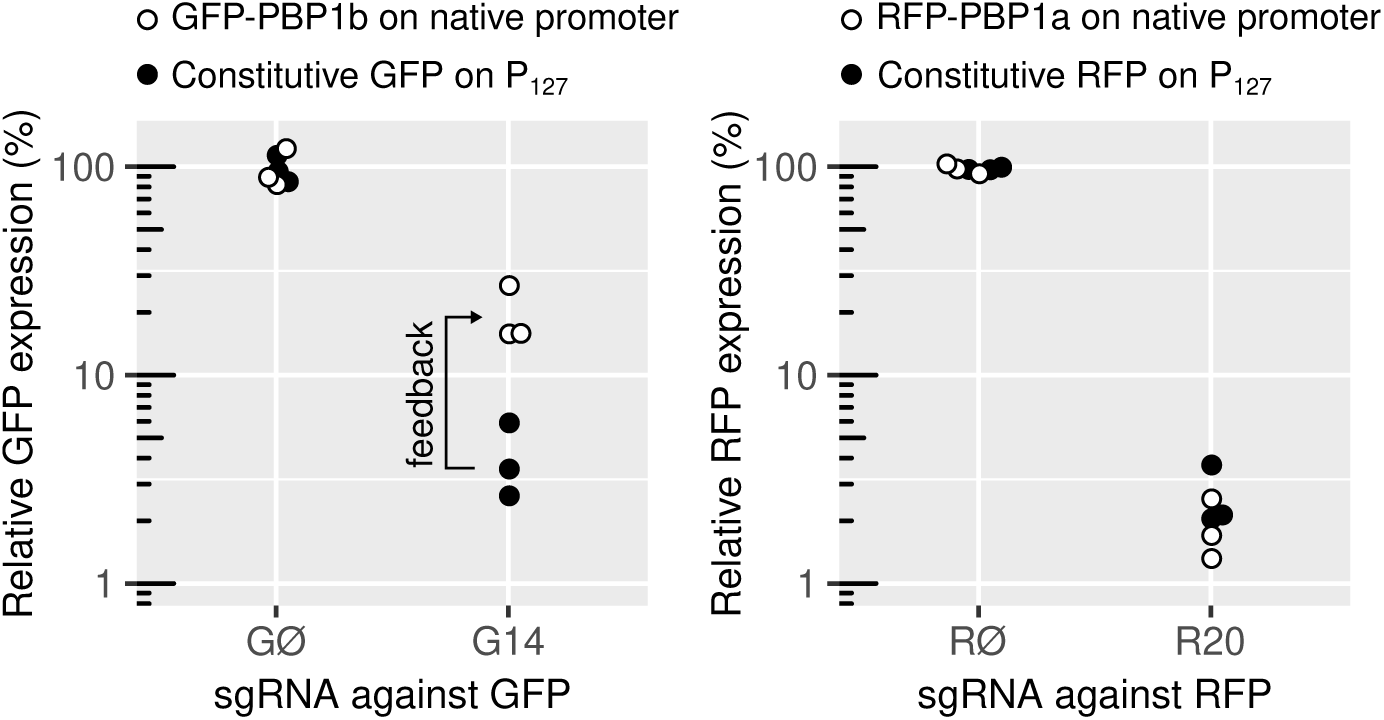
Residual PBP1a and PBP1b levels in response to CRISPR-based repression, measured by fluorescence microscopy. **Left:** The same CRISPR guides produce different repression strength on GFP, depending on whether it is expressed constitutively in AV47 (186::P_Tet75_-dCas9, HK022::P_127_-*sfgfp*, λ::P_127_-*mcherry*) or fused to PBP1b in the native locus (AV44). **Right:** Same experiment on the RFP-PBP1a fusion, on AV44, showing no evidence for feedback. sgRNAs are expressed from pAV20 with sgRNA GØ, RØ, G14 or R18 as annotated. In each case, relative fluorescence is expressed as a percentage of the fluorescence of the same strain carrying the pAV20 GØ-RØ control plasmid, i.e. without repression.

**Figure 1 - Supplement 5.**
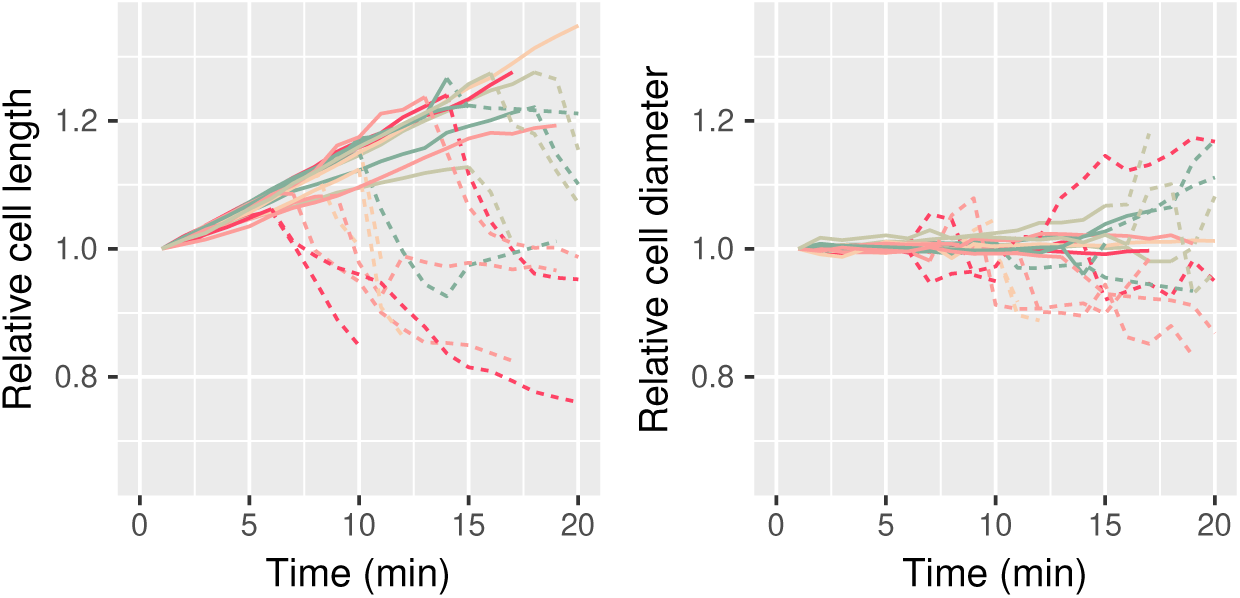
Dimensions of individual cells before lysis due to PBP1ab repression. Strain: AV44/pAV20 G20-R20. Imaging starts 4h45 after the induction of the CRISPR system. Cell length and cell diameter are normalized with respect to the dimensions of the cell in the first frame of the movie. Solid lines are living cells, dashed lines are lysed cells (phase-bright). Colors are arbitrary.

**Figure 1 - Supplement 6.**
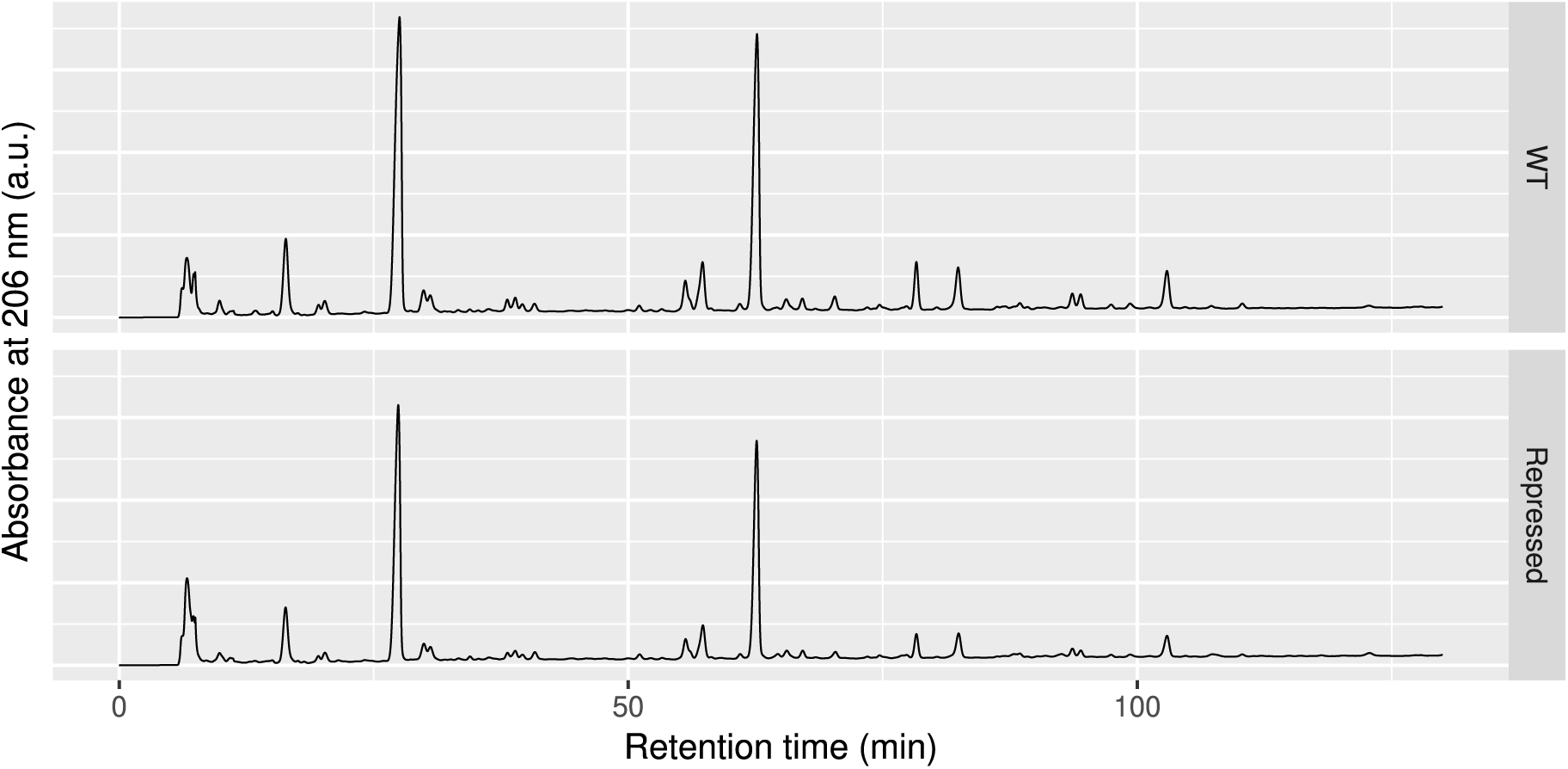
HPLC analysis of the peptidoglycan after digestion by mutanolysin in LC69 (WT) and AV44/pAV20 G14-R20 (Repressed). a.u.: arbitrary units.

**Figure 2 - Supplement 1.**
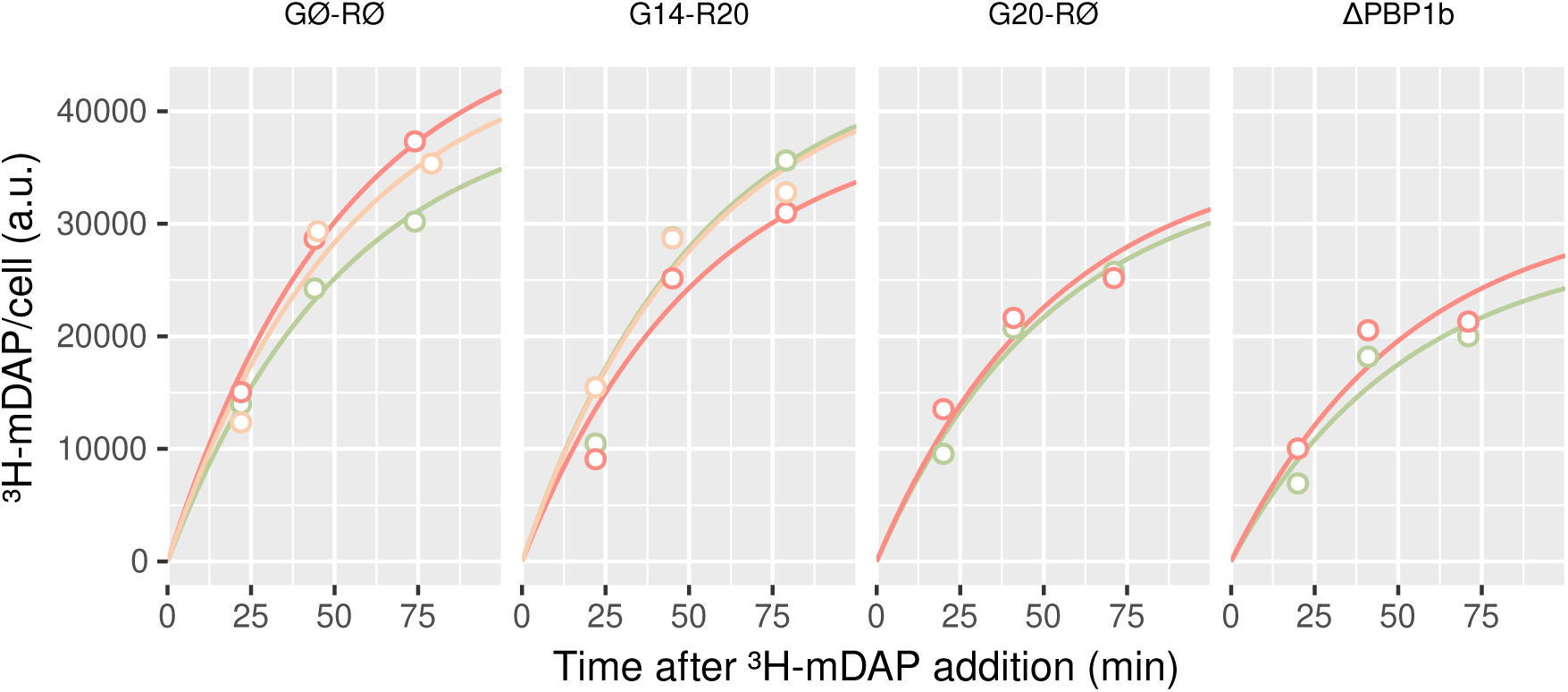
Amount of incorporated ^3^H-mDAP per optical density as a function of time. Strains are AV84 (AV44 Δ*lysA*)/pAV20 GØ-RØ, AV84/pAV20 G14-R20, AV84/pAV20 G20-RØ and AV105 (AV84 ΔPBP1b)/pAV20 GØ-RØ, from left to right. Colored curves are exponential fits (see methods) to the raw measurements (open symbols). Each color represents one biological replicate. a.u.: arbitrary units, corresponding to CPM per optical density units.

**Figure 2 - Supplement 2.**
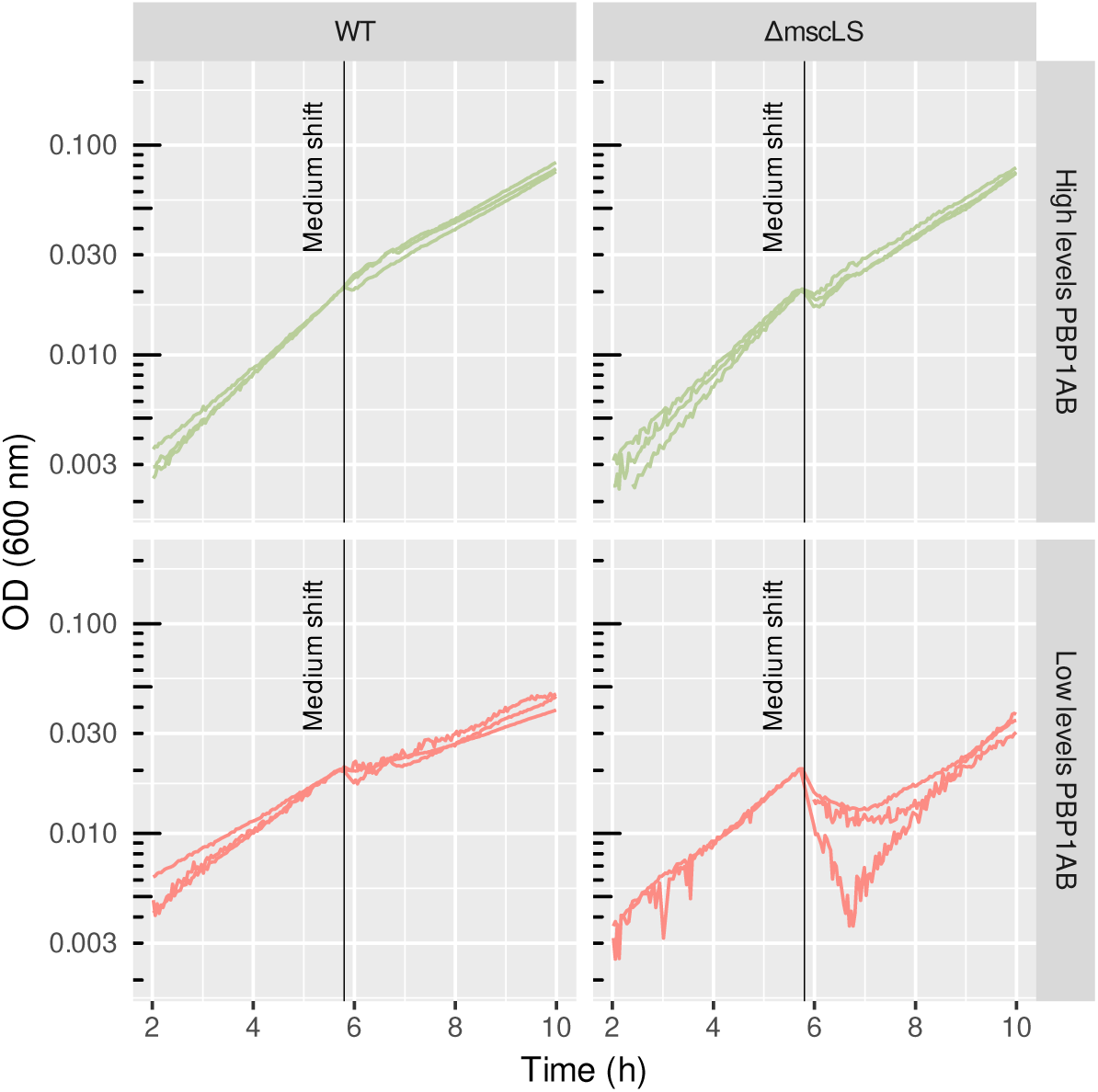
Growth curves before and after osmotic shock. Strains are AV44 (“WT”) or AV93 (AV44 Δ*mscSL*) (“ΔmscSL”) with pAV20 GØ-RØ (“High levels PBP1ab”) or pAV20 G14-R20 (“Low levels PBP1ab”). Vertical lines mark the time of centrifugation, medium removal and resuspension in a medium of lower osmolarity. Optical densities are normalized with respect to the value at the time of the medium shift. Each curve is one biological replicate. Experiment done in a plate-reader. OD: Optical density.

**Figure 3 - Supplement 1.**
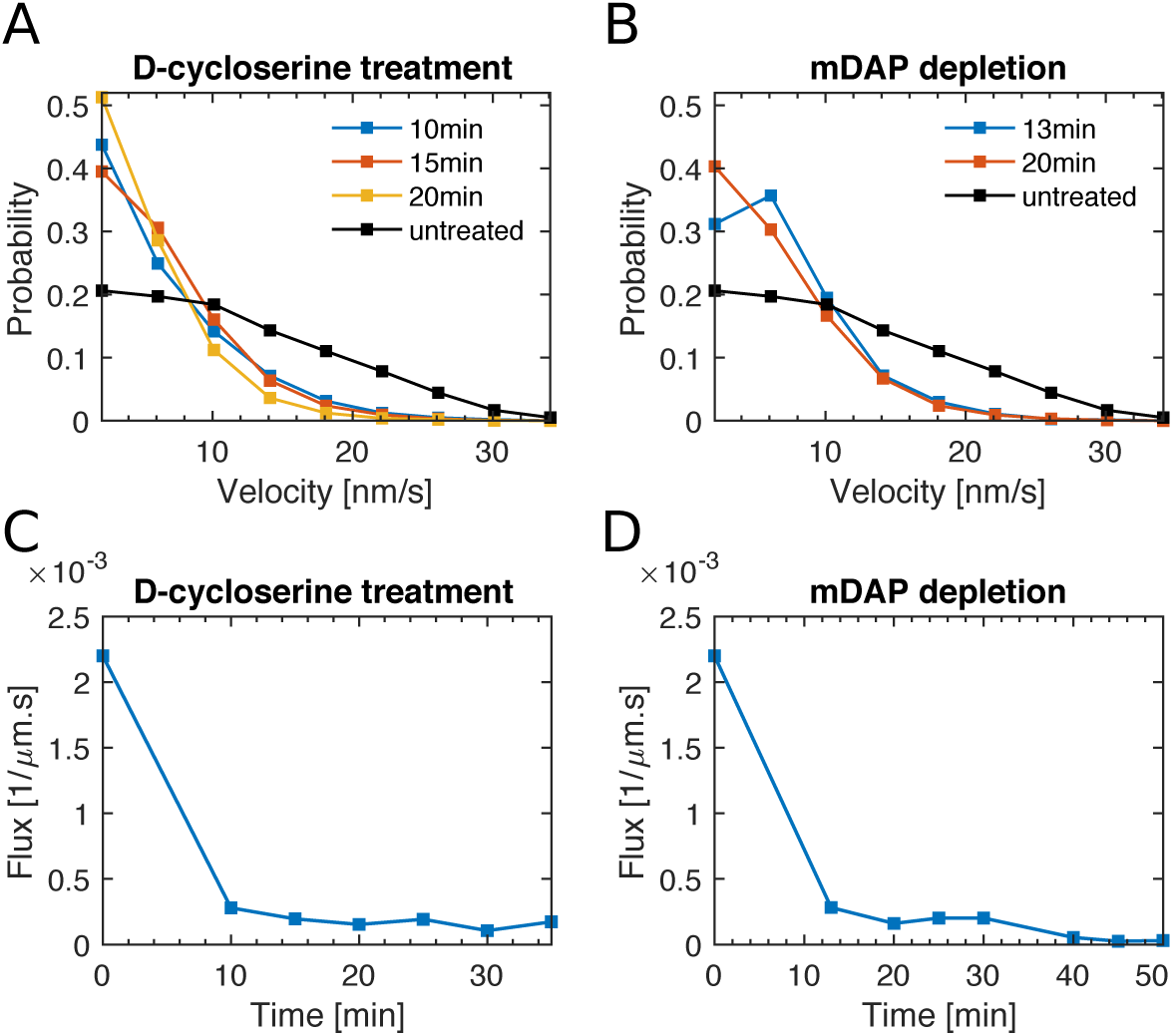
Effect of depletion of peptidoglycan precursors measured by MreB motion. **(A, B):** Probability distributions of the instantaneous velocity of MreB-msfGFP measured upon D-cycloserine treatment using strain B172 (MG1655 mreB<>mreB-msfGFP) (A), or during depletion of mDAP in in mDAP auxotroph, B176 (MG1655 asd-1, mreB<>mreB-msfGFP) (C). Measurements carried out in LB. **(B, C):** Flux of MreB-msfGFP corresponding to the experiments in A, B, respectively. Flux is calculated as sum of track end-to-end distances divided by total cell area and movie duration.

**Figure 3 - Supplement 2.**
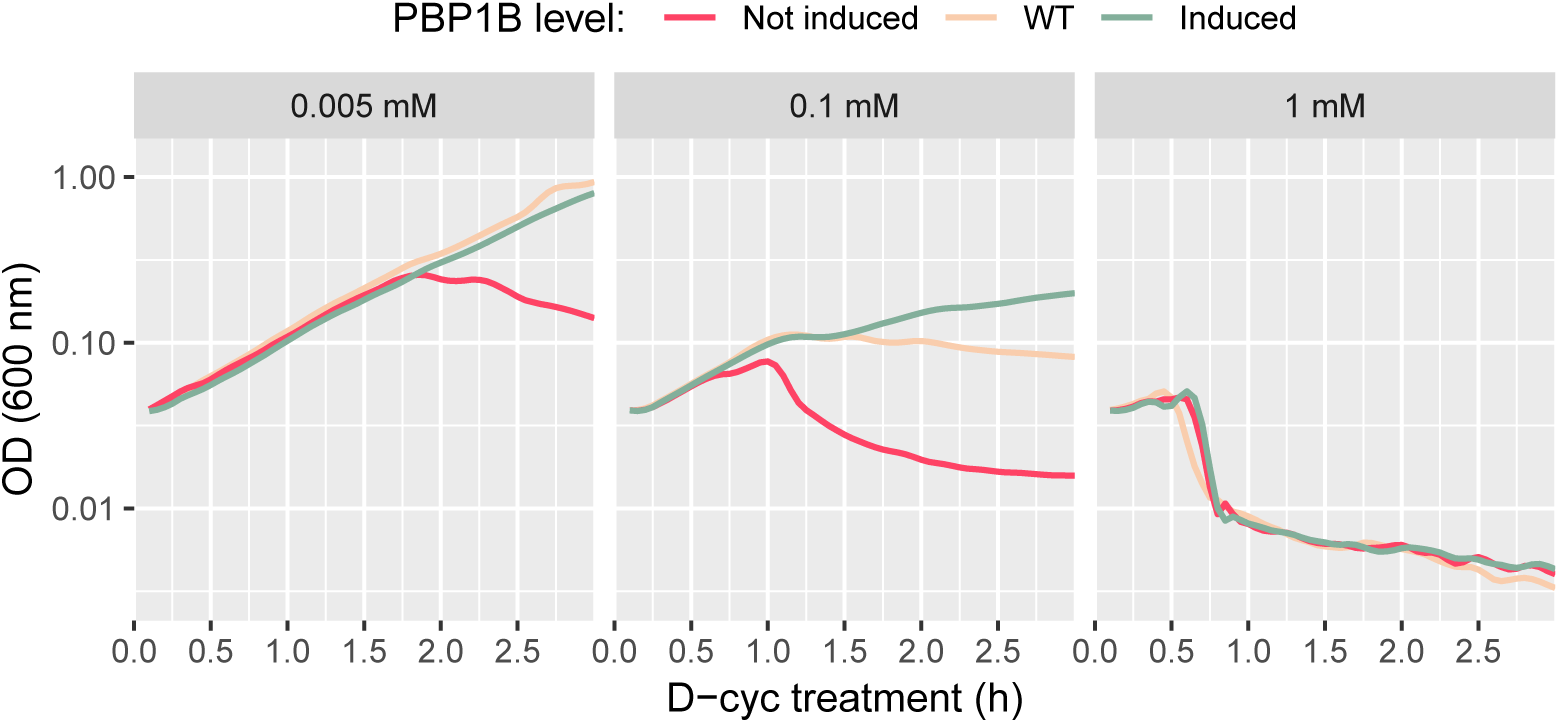
D-cycloserine sensitivity at different PBP1b levels and drug concentrations. Growth curves of B150 (ΔPBP1b)/pBC03 (pBAD33-P_Bad_-PBP1b) induced with arabinose (“Induced”), not induced (“Not induced”) or MG1655 (“WT”), during treatment with three different concentrations of D-cycloserine (0.005 mM to 1 mM). Measurements performed in plate reader. OD: Optical density.

**Figure 4 - Supplement 1.**
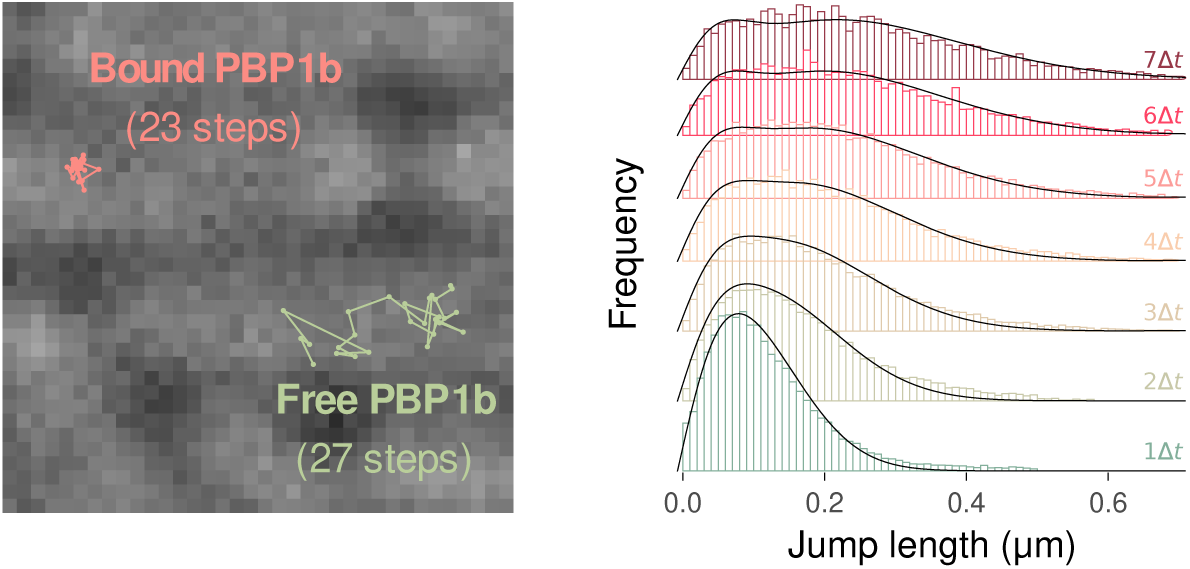
Tracking of single molecules using the Spot-On tool. **Left:** Sample tracks corresponding to bound and diffusive GFP-PBP1b molecules, overlaid on a brightfield image using strain AV44/pCRRNAcos G10-R18 (280% PBP1a, 130% PBP1b). **Right:** Observed and fit distributions of particle jump lengths over *n* time steps. Distributions shown for one replicate of strain AV44/pCRRNAcos G10-R18.

**Figure 4 - Supplement 2.**
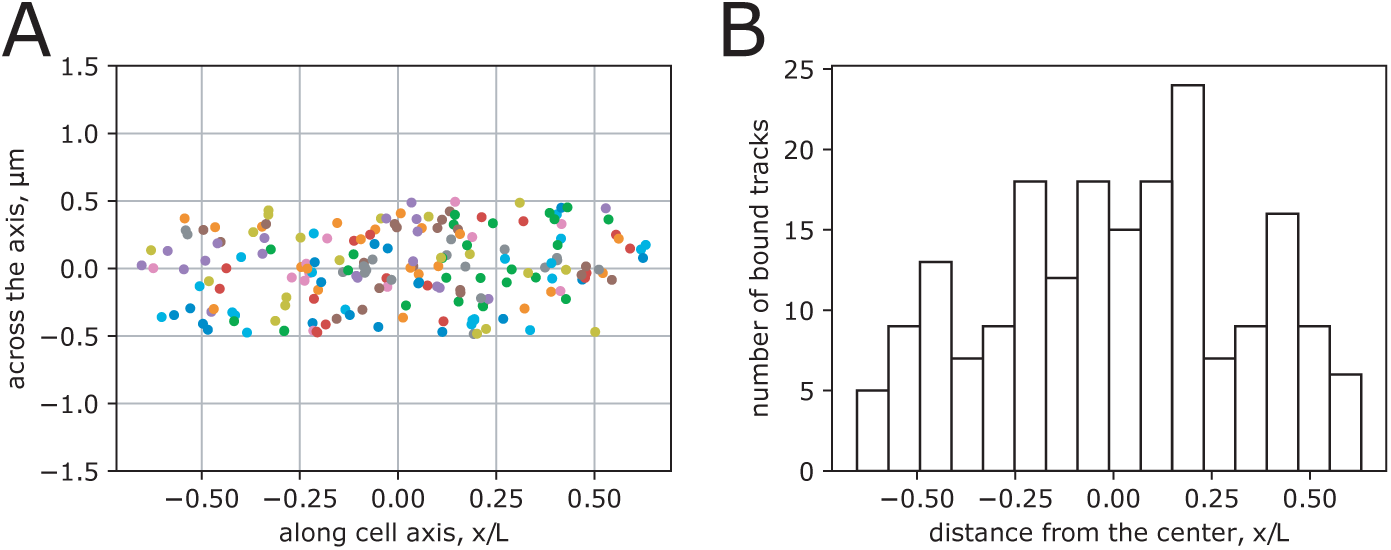
Localization of bound molecules. **(A-B):** Position of bound molecules with respect to a normalized coordinate system measured in AV51 (AV44 ΔPBP1a)/pCRRNAcos with crRNA G10 (130% PBP1b with respect to WT). Bound molecules were identified according to their MSD (MSD *<* (50nm)^2^. In 67 cells we found 195 tracks. Tracks were assigned to a cell if the distance from the axis of the cell was below 0.5 μm. The average length of cells was 3.7 μm. (A) scatter plot of bound sites with normalized longitudinal coordinate. (B) histogram along x of normalized cell.

**Figure 4 - Supplement 3.**
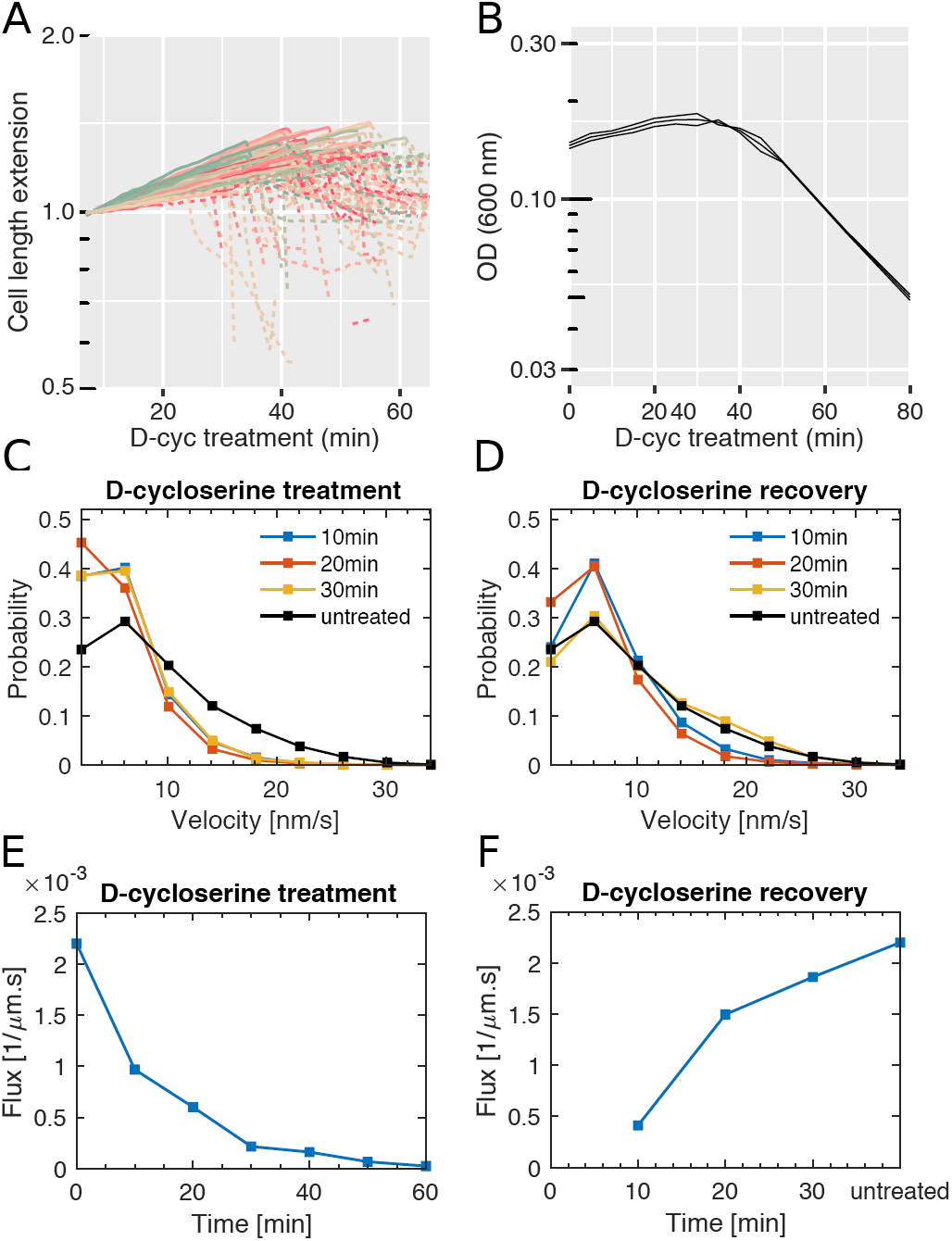
Effect of D-cycloserine treatment in minimal media. **A:** Cell length during D-cycloserine treatment (1 mM) under the microscope using strain AV51 (AV44 ΔPBP1a)/pAV20 in minimal medium with glucose and 0.01% casamino acids.Length is normalized by the length of the same cell at *t* = 0. Solid lines: growing cells, dashed lines: phase-bright, lysing cells. **B:** Growth curves of AV51 during D-cycloserine treatment (1 mM; same medium as in A).Lines represent three biological replicates. OD: Optical density. **C-F:** MreB rotation during D-cycloserine treatment (C,E) and during recovery from a 30 min period of D-cycloserine treatment (D,F) using strain B172 (MG1655 mreB<>mreB-msfGFP) grown in minimal medium as in A and analyzed as in Figure 3 - Supplement 1.

**Figure 4 - Supplement 4.**
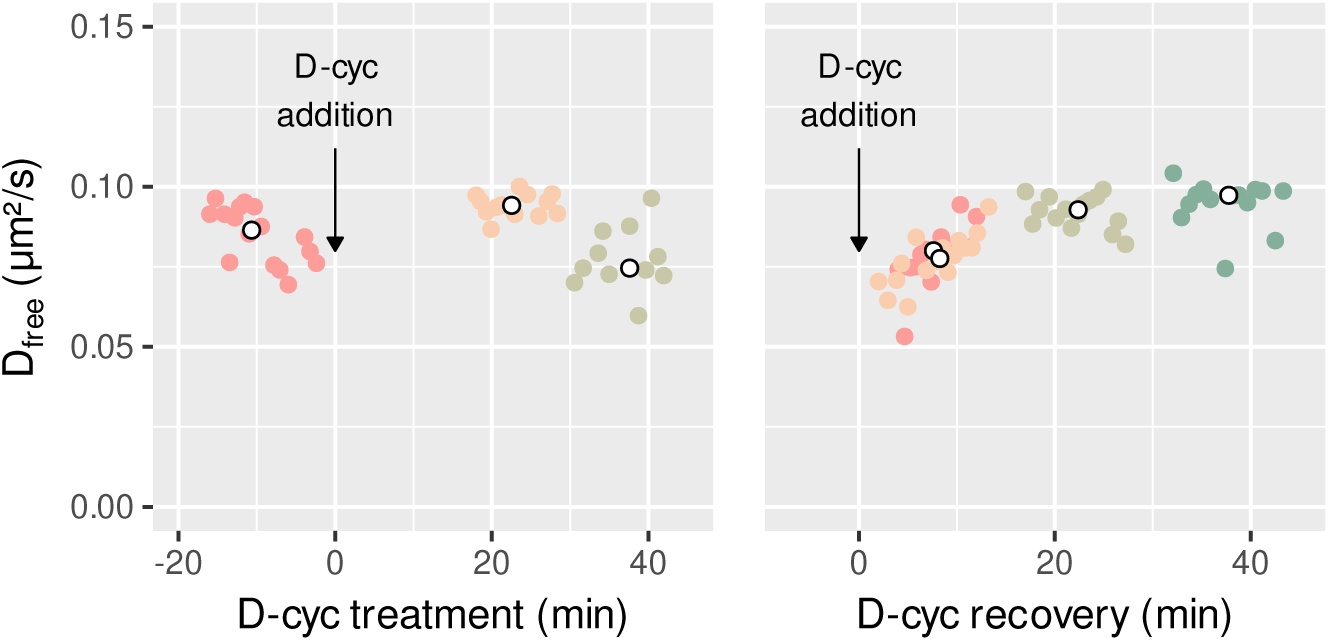
Free diffusion coefficient (*D*_free_) during D-cycloserine treatment and recovery. **(A-B):** *D*_free_ of GFP-PBP1b at different times during 1mM D-cycloserine treatment (A) and during recovery from 30 min of 1 mM D-cycloserine treatment (B) in the strain AV51/pCRRNAcos G10-RØ. Corresponding bound fractions provided in Figure 4 (D-E).

## Supplementary information

### Quantification of GFP-PBP1b by SDS-page

A sfGFP-His6 fusion protein was purified in this study to be used as an internal standard for the semi-quantitative sfGFP-PBP1b SDS-page. The sfGFP-6xHis fusion was expressed and purified from a BL21(DE3) *E. coli* strain. A 10 ml LB preculture containing carbenicillin (100 μg/ml) was inoculated from a freshly transformed colony and grown at 37°C until an OD_600_≈0.6. This culture was diluted 1:100 into 500 ml fresh pre-warmed LB containing carbenicillin (100 μg/ml) and grown at 37°C to an OD_600_≈0.6. At this time point, the expression was induced by the addition of 1 mM of IPTG and the culture was incubated at 20°C overnight. The next day, the culture was cooled for 15 min at 4°C and the cells were recovered by centrifugation (4,000 x g) at 4°C for 15 min. Cell pellets were resuspended in 12,5 ml of lysis buffer (20 mM Tris-HCl pH 8, 100 mM NaCl, 5 mM 2-mercaptoethanol, 20 mM imidazole, 1 mM PMSF) and stored at −80°C. Cells were thawed, benzonase (E1014, Millipore) and lysozyme (L6876, Sigma) were added (respectively 500 units and 0,5 mg/ml) and cells were disrupted by sonication on ice. Cell debris and membranes were pelleted by centrifugation at 40,000 × g for 1 hour at 4°C. In parallel, a 2 ml aliquot of Ni-NTA agarose resine slurry (#25214, Thermoscientific), corresponding to a 1 ml beads volume, was equilibrated using 50 ml of buffer with 20 mM of imidazole. The soluble protein extract was incubated with the beads for 1 hour on a wheel at 4°C and loaded on a gravity column. The beads were extensively washed on the column using 50 ml of buffer (20 mM Tris-HCl pH 8, 500 mM NaCl, 5 mM 2-mercaptoethanol, 50 mM imidazole, 10% glycerol). Bound sfGFP-6xHis proteins were eluted in 10 ml buffer (20 mM Tris-HCl pH 8, 500 mM NaCl, 5 mM 2-mercaptoethanol, 120 mM imidazole, 10% glycerol). Fractions of 1 ml were collected and their concentration was estimated using a Bradford-based Protein Assay (Bio-Rad) according to the instructions. The purity of elution fractions was also estimated by loading 5 μl on a 4-20 percent polyacrylamide gel (Miniprotean TGX, Bio-rad) stained with Coomassie blue and scanned with a Typhoon 9000 FLA imager (GE Healthcare) to detect GFP signal (473 nm laser, excitation wavelengh 489 nm, emission 508 nm) (figure 1 - Supplement 3A).

In order to estimate the copy numbers of sfGFP-PBP1b per cell, three independent cell extract preparations of AV44 pAV20-GØ-RØ (non-repressed), AV44 pAV20 G14-R20 and AV51 (ΔPBP1a) pAV20 G14-R20 were analyzed by fluorescence gel-based assay. Cells were grown overnight in LB at 30°C and diluted 1/100 into 40 ml of LB with 100 ng/ml anhydrotetracycline. Three independent cultures, for each strain, were grown at 30°C to an OD_600_ approximately of 0.3 and the colony forming units (cfu) of each culture were determined by plating serial dilutions on LB plates. Cells were harvested by centrifugation, resuspended in 200 μl of PBS 1x. Cells were disrupted by sonication and protein concentrations were determined using a Bradford-based Protein Assay (5000006, Bio-Rad) according to the instructions. 150 μl of the total cell extract was mixed with 25 μl of Laemmli sample buffer 4X (#1610747, Bio-Rad). Cell extracts were flash-freezed in liquid nitrogen and stored at −80°C. To determine the amount of PBP1b in each of the extracts, normalized amounts of total protein were loaded on 4-20 % polyacrylamide gels (Miniprotean TGX, Bio-rad) together with increasing amounts of purified sfGFP-6xHis (same sfGFP used for PBP1b tagging). After migration, the gel was stained with Coomassie blue and scanned for fluorescence as detailed above. A standard curve plotting integrated signal intensity versus protein concentration was generated for the purified sfGFP-6xHis and was used to determine the number of molecules of sfGFP-PBP1b loaded on the gel for each cell extract. The cell number determined for the initial cell cultures were then used to calculate the number of PBP1b molecules per cell.

